# Avian reovirus triggered ferroptosis, modulated lipid metabolism, and induced lipid droplet formation in infected cells, which was mediated in part by the p10/FAST protein

**DOI:** 10.64898/2026.11.10.765664

**Authors:** Megan Liu, Declan Kehlbeck, Sofia Egana-Labrin, Andrew Brodrick, Leo Sun, Jace Jones, Nishanth Sunny, Andrew J. Broadbent

## Abstract

In this study, we evaluated how avian reovirus (ARV) affected lipid metabolism and lipid droplets (LDs) in cell culture. ARV infection of chicken LMH cells significantly increased the size and number of LDs, however, LDs did not co-localize with virus factories, and neither overexpression by treatment with fatty acids (FAs), nor inhibition by treatment with TOFA or DGAT inhibitors significantly affected ARV replication. Lipidomic analysis revealed that following infection, total cellular lipid content increased approximately 25-fold, phosphatidylcholine (PC) content increased from 23.5 to 57.9%, and the polyunsaturated FA:monounsaturated FA ratio (PUFA:MUFA) decreased in membrane phospholipids but increased in triglycerides. Transcriptomic analysis revealed that there was also a reduction in the expression of genes involved in PUFA synthesis. As membrane PUFAs are susceptible to peroxidation, we investigated whether ARV induced ferroptosis. Consistent with this, ARV led to an increase in oxidative stress and lipid peroxidation that was partially iron dependent, and we found evidence that ferroptosis was proviral. Moreover, as ARV is known to form syncytia, we hypothesized that our observations were also the result of plasma membrane remodeling. Consistent with this, cells transfected with the ARV fusogenic p10/FAST protein formed syncytia, had a significantly upregulated number of LDs, and displayed hallmarks of ferroptosis. Taken together, we hypothesize that ARV induces syncytium formation and ferroptosis to facilitate viral spread to neighboring cells, but to prevent premature cell death due to these processes, lipid metabolism is modulated by infection, leading to an increase in LDs.

**Importance:** Lipid droplets (LD) are utilized by some reoviruses to facilitate replication. However, in this study, we found that although avian reovirus (ARV) induced LD formation in infected cells, LDs did not play a significant role in viral replication. Instead, we demonstrated that, in addition to inducing syncytia, ARV triggered ferroptosis and modulated lipid metabolism in infected cells. Moreover, we found that the p10-FAST protein contributed to these processes. We propose that ARV-induced syncytia and ferroptosis facilitate viral spread to neighboring cells, but to prevent premature cell death due to these processes, lipid metabolism is modulated by infection, leading to an increase in LDs as a byproduct. Reoviruses are known to restrain cell death to favor replication. Here, our data suggest that lipid metabolism may be modulated to achieve this goal.

## Introduction

Cellular lipid droplets (LDs) are small cytoplasmic organelles comprised of a neutral lipid core containing triglycerides (TGs) and sterol esters, surrounded by a phospholipid monolayer and LD-associated proteins^1^. LDs are involved in the regulation of numerous cellular processes, such as endoplasmic reticulum (ER) stress, oxidative stress, autophagy, and energy homeostasis^1,2^. Several viruses have been reported to utilize LDs to promote their replication within cells, including double-stranded (ds) RNA viruses in the family *Reoviridae*^3–7^. Within the *Reoviridae* family, Rotaviruses (RVs) were found to utilize LDs as a scaffold on which viral replicative complexes, known as viroplasms, form. RV infection induced LD formation, and viral nonstructural proteins NSP2 and NSP5, which are involved in viroplasm formation, colocalized with LD-associated proteins Perilipin (PLIN) 1 and PLIN2^3,7^. Moreover, pharmacological inhibition of the *de novo* lipogenesis pathway by treatment with 5-(tetradecyloxy)-2-furancarboxylic acid (TOFA) and/or C75 decreased LD formation, viroplasm formation, viral RNA replication, and infectious titer^3,4,8^. In addition, Grass carp reovirus (GCRV) infection also led to the upregulation of LD formation in infected cells, and inhibition of LD formation by treatment of cells with TOFA decreased GCRV replication, whereas upregulation of LD formation by treatment with palmitic acid increased replication^5^. Furthermore, the GCRV protein NS80, which drives viroplasm formation, colocalized with LDs, consistent with the virus utilizing LDs to form its replicative compartments. In contrast, mammalian orthoreovirus has been shown to utilize LDs for temporospatial orchestration of viral genome amplification and virion assembly by binding outer capsid proteins, and redirecting them away from less mature replicative complexes, known as virus factories, toward more mature factories in the perinuclear region, to prevent premature termination of viral genome amplification by outer capsid assembly onto core particles^6^.

Despite this growing body of knowledge, little is known about the role LDs play in the replication cycles of avian dsRNA viruses. In this study, we investigated the role of LDs in the context of avian reovirus (ARV), a member of the *Reoviridae* family, and infectious bursal disease virus (IBDV), a member of the *Birnaviridae* family, both of which form VFs in infected cells. As previous research has shown that LD induction in infected cells is pro-viral for dsRNA viruses, we hypothesized that ARV and IBDV infection would lead to upregulation of LD formation, and that LD upregulation would be pro-viral and increase viral replication. To test these hypotheses, we aimed to establish whether: 1) ARV and IBDV infection increased LD number, 2) viral proteins colocalized with LDs, and 3) inhibition or upregulation of LD synthesis affected viral replication. Furthermore, research has shown that modulation of lipid metabolism, which can result in the formation of LDs, is critical in regulating redox homeostasis and ferroptosis sensitivity during cell stress, such as during viral infection^2,9^. Therefore, we explored how infection influenced lipid metabolism and ferroptosis.

## Results

### ARV infection induced the formation of LDs, but IBDV had a less significant effect

Infection of the chicken hepatic cell line, LMH, with ARV strain S1133 significantly increased LD number 6-24 hours post-infection (hpi), while also increasing LD size, most notably at 24hpi, using LipidTOX Red as a marker for LDs (Figure 1, A-C). In contrast, infection with IBDV strain PBG98 had a less pronounced effect on LD formation (Figure 1, D-F). We then repeated the experiment in DF-1 cells, which are more susceptible to IBDV infection compared to LMH cells, and in chicken primary gut cultures, to investigate any differences between immortalized cell lines and primary cells. Consistent with results in LMH cells, ARV infection increased LD number in DF-1 cells, while IBDV did not (Figure S1, A-C), and ARV also increased LD number in primary chicken gut cultures (Figure S1, D-F). Furthermore, chicken field strains of ARV (GF2-E and B8-T) also led to an increased number of LDs in LMH cells (Figure S2, A-C), demonstrating that the phenomenon was not limited to one strain. Taken together, ARV infection increased LD formation across various cell types and viral strains, while IBDV had a less significant effect, suggesting that the two viruses had different effects on lipid metabolism in infected cells.

**Figure 1.**
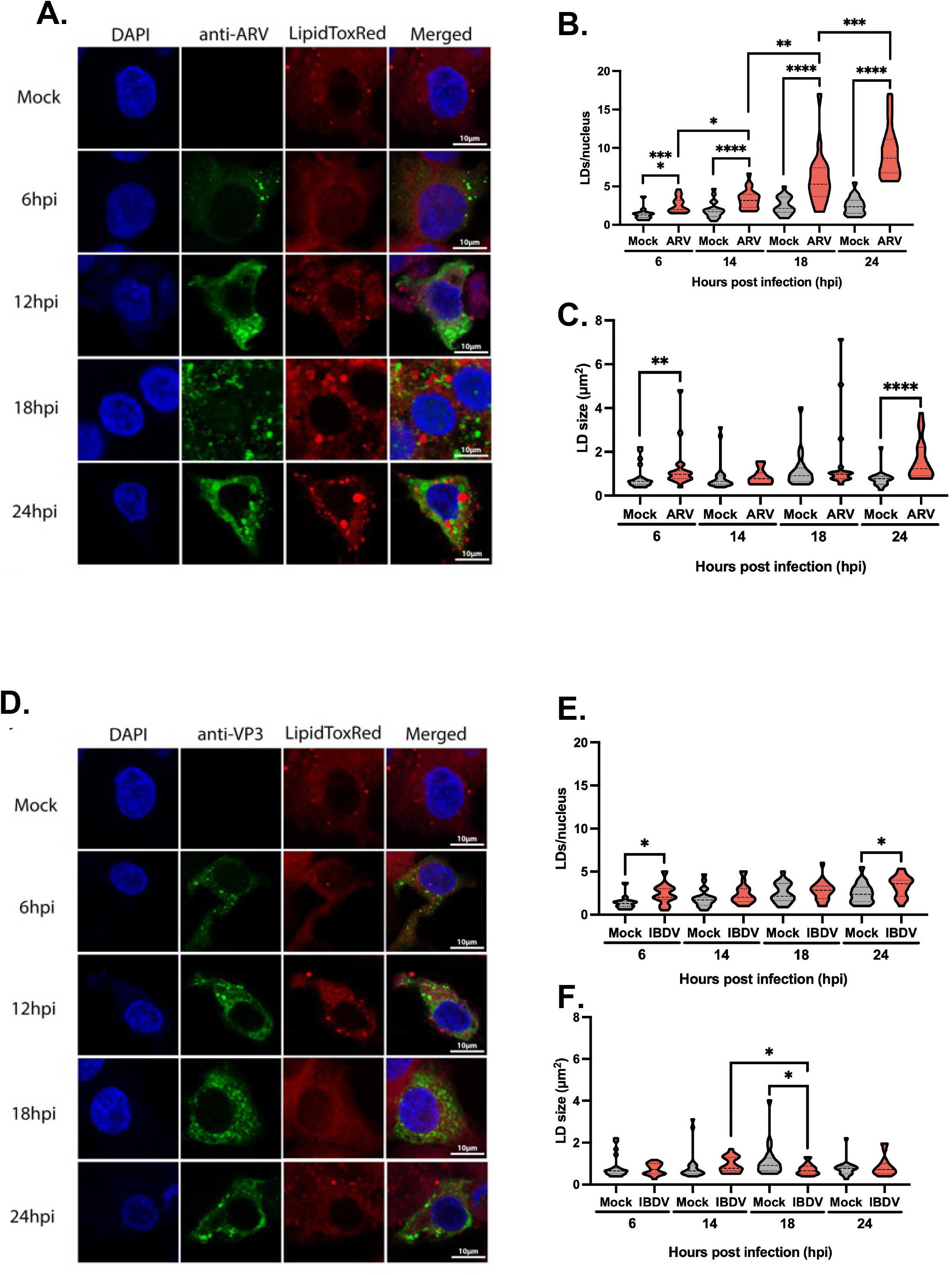
ARV infection induced the formation of LDs while IBDV had a less significant effect. LMH cells were seeded onto coverslips and infected with ARV S1133 (A-C) or IBDV PBG98 (D-F) at an MOI of 1. At each timepoint, cells were fixed with 4% PFA and stained with ARV polyclonal antibody (A-C) or VP3 antibody (D-F) and LipidTOX Red. Cells were imaged and LD number per nucleus and LD area were quantified using a scikit-image segmentation pipeline. The Mann-Whitney U Test was used for statistical analysis in GraphPad Prism. \**p* < 0.05, \*\**p* < 0.01, \*\*\**p* < 0.001, \*\*\*\**p* < 0.0001.

### LDs did not colocalize with ARV VFs, and neither inhibition nor promotion of LDs had a significant effect on viral replication

ARV-infected cells contained viral cytoplasmic puncta that did not show a strong colocalization with LDs (Figure 1A), suggesting that LDs did not colocalize with ARV VFs. However, we used a polyclonal anti-ARV antibody to visualize infection, which bound multiple viral proteins, and, as such, it was not possible to confirm that the puncta were true VFs. The non-structural protein µNS is a critical component of ARV VFs^10,11^. Therefore, we transfected LMH cells with a plasmid expressing mNeonGreen (mNG) tagged to the N-terminus of µNS from ARV strain S1133, to induce VF formation that we could visualize. When cells were transfected with the mNG:: µNS -expressing plasmid alone, or co-transfected and infected with ARV S1133, cytoplasmic puncta were visible by fluorescence microscopy. ARV VFs did not colocalize with LDs, unlike what is reported for RV and GCRV, suggesting that ARV does not utilize LDs as a scaffold for VF formation (Figure 2).

**Figure 2.**
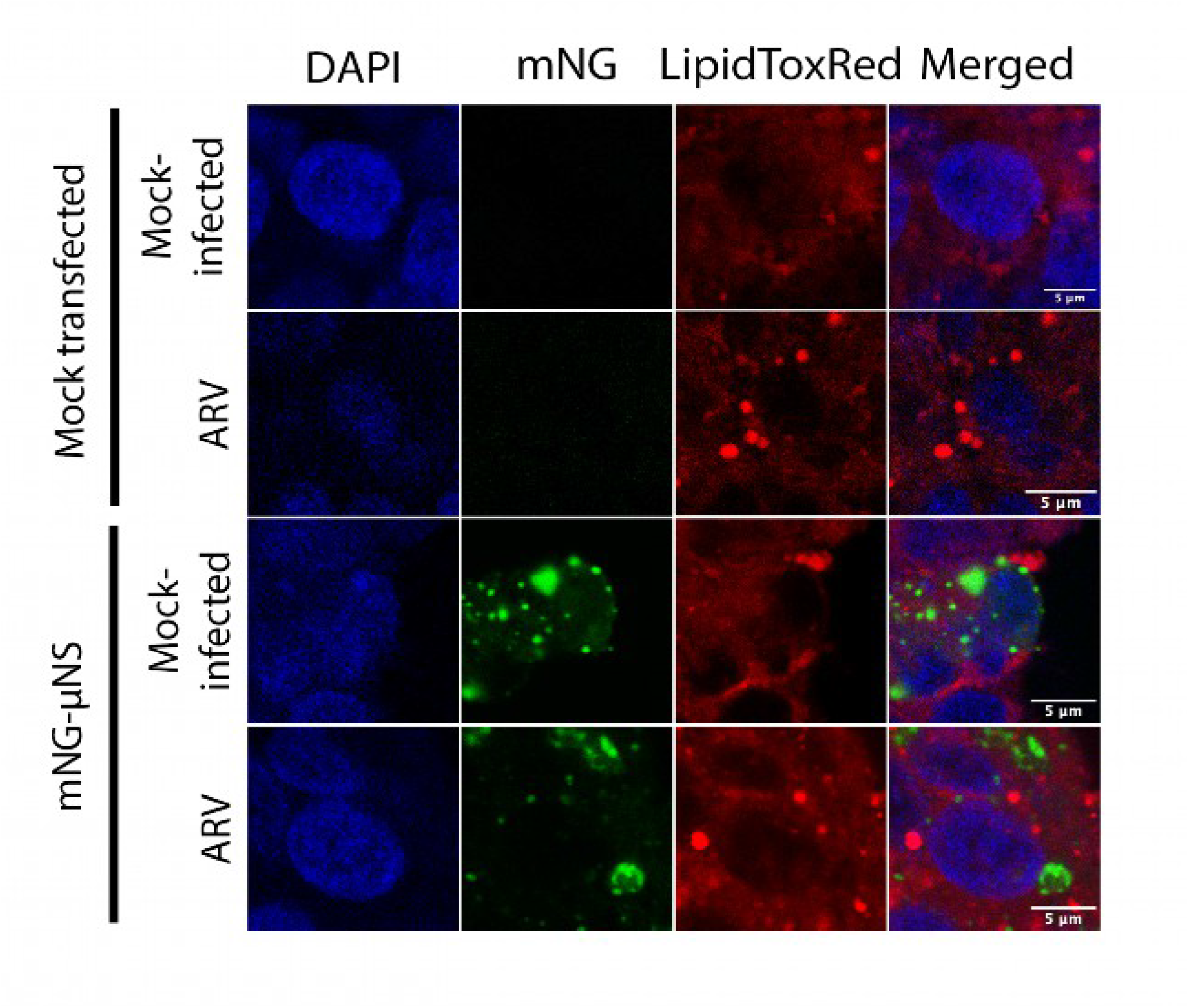
ARV VFs did not colocalize with LDs. LMH cells were transfected with a CMV promoter-driven mNG::µNS construct and co-infected with ARV S1133 at an MOI of 1. Mock transfection control cells received an empty vector. Cells were fixed with 4% PFA at 24 hours post-transfection, stained with LipidTOX Red, and imaged to evaluate colocalization between µNS positive VFs and LDs.

Previous research showed that induction of LDs during RV or GCRV infection is proviral, as inhibition of LD formation decreased viroplasm formation and viral replication, while promotion of LD formation had the opposite effect^3,5,8^. To determine whether LD induction was proviral during ARV or IBDV infection, we inhibited LD formation with TOFA, an allosteric inhibitor of acetyl-CoA carboxylase 1 (ACC1), which catalyzes the first, rate-limiting step of *de novo* lipogenesis and is therefore essential for LD formation^8^. We performed MTT cell viability assays with varying concentrations of TOFA and determined that a working concentration of 15µM did not induce cytotoxic effects in LMH cells (Figure S3, A). TOFA treatment alone decreased LD number, but ARV infection still resulted in LD upregulation (Figure 3 and Figure S3, B). We hypothesized that ARV-infected cells could still acquire free FAs exogenously from the FBS in the cell culture media despite inhibition of intracellular *de novo* lipogenesis by TOFA. Therefore, LMH cells were cultured in FA-free conditions in media without FBS and instead supplemented with 1mg/mL FA-free BSA. TOFA treatment combined with FA-free culture conditions abrogated LD upregulation upon ARV infection (Figure 3 and Figure S3, B), but there was no effect on viral replication measured by either RT-qPCR or tissue culture infectious dose-50 (TCID_50_) for either ARV or IBDV (Figure 3).

**Figure 3.**
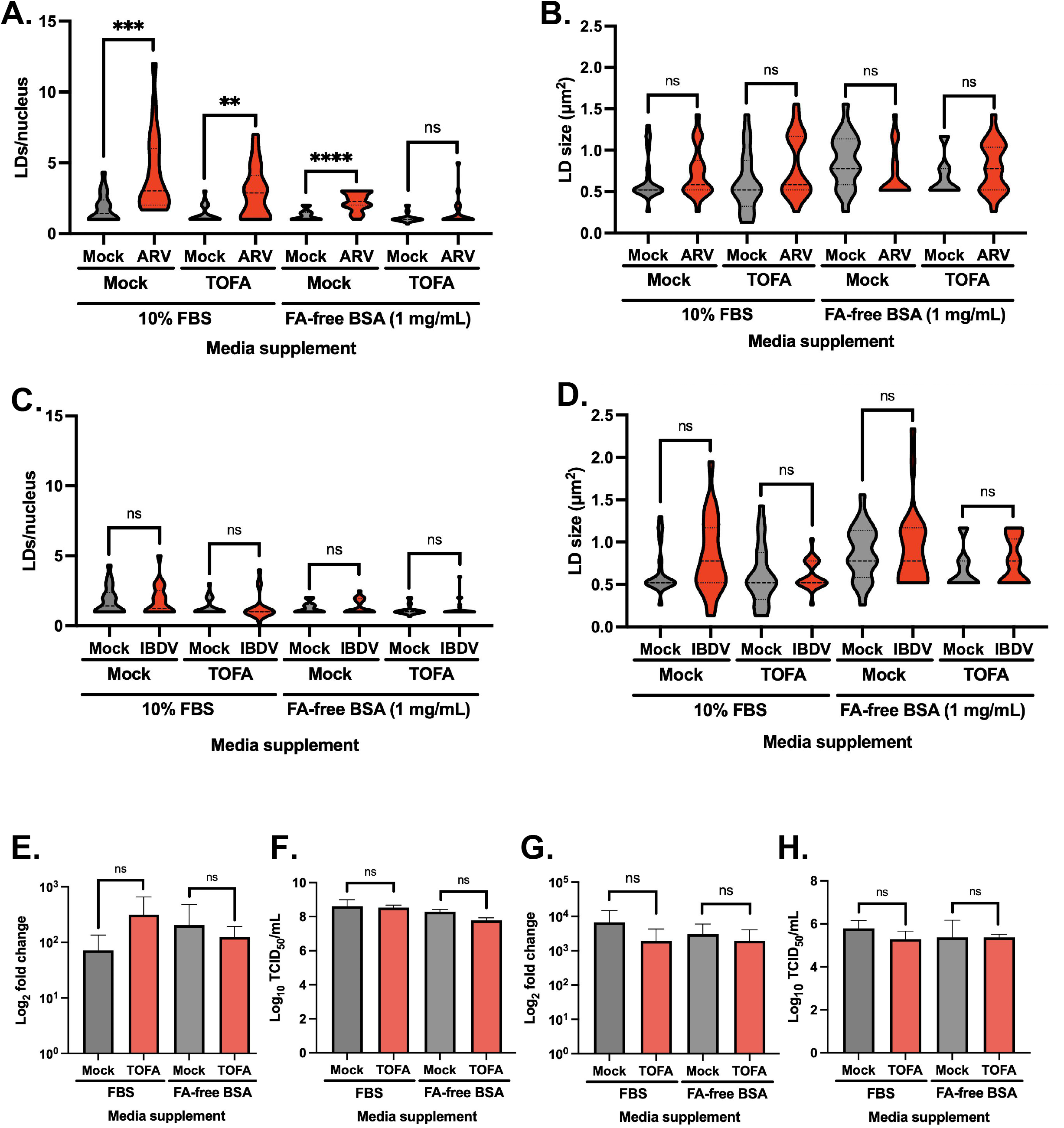
Inhibition of LD formation with TOFA had no effect on ARV or IBDV viral replication. LMH cells were cultured in either DMEM F-12 supplemented with 10% FBS or in DMEM F-12 with 1mg/mL FA-free BSA for one passage before seeding onto 24 well plates. Cells were pre-treated with 15µM of TOFA 8 hours before infection with ARV S1133 (A, B, E, and F) or IBDV PBG98 (C, D, G, and H) at an MOI of 1. At 18hpi, cells were fixed in 4% PFA for LipidTOX Red staining and LD quantification (A-D), lysed in RLT buffer for RNA extraction and viral transcript RT-qPCR (E, G), or culture medium supernatant were collected for TCID_50_ (F, H). \**p* < 0.05, \*\**p* < 0.01, \*\*\**p* < 0.001, \*\*\*\**p* < 0.0001.

Conversely, we induced LD formation through FA supplementation by treating LMH cells with palmitic acid (PA) and/or oleic acid (OA) conjugated to BSA. An MTT assay revealed no difference in cell viability following treatment (Figure S4, A). Treatment resulted in increased LD number and size, but there was no significant effect on ARV or IBDV viral replication (Figure 4, and Figure S4). Taken together, these data demonstrate that LDs are not proviral for either IBDV or ARV, suggesting that LDs play a less significant role in ARV VF formation and replication than they do in RV or GCRV infection.

**Figure 4.**
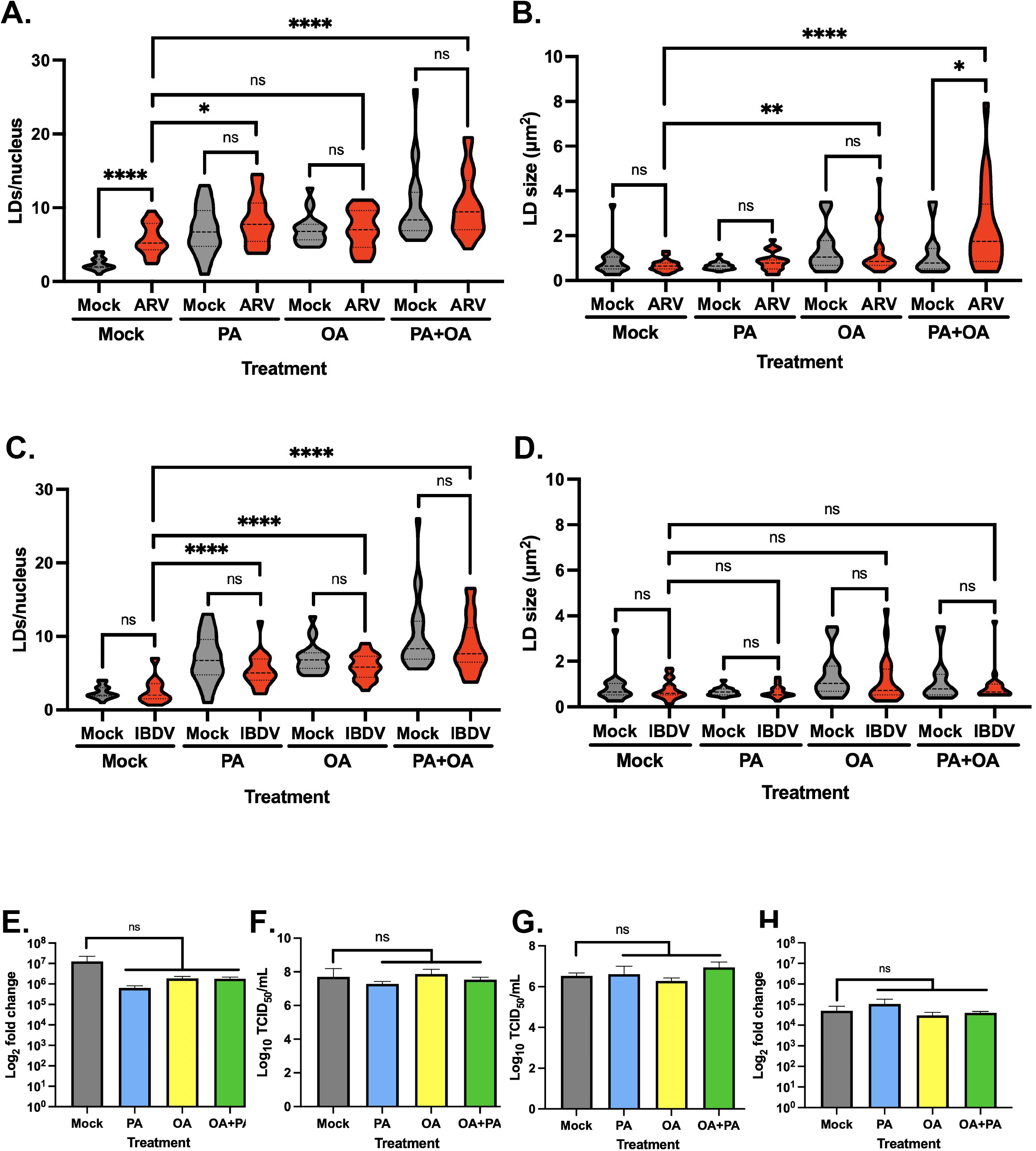
Induction of LD formation with exogenous fatty acid supplementation had no effect on ARV or IBDV viral replication. LMH cells were pre-treated with PA-BSA, OA-BSA, or a 2:1 mixture of PA-BSA/OA-BSA 8 hours before infection with ARV S1133 (A, B, E, and F) or IBDV PBG98 (C, D, G, and H). At 18hpi, cells were fixed in 4% PFA for LipidTOX Red staining and LD quantification (A-D), lysed in RLT buffer for RNA extraction and viral transcript RT-qPCR (E, G), or culture medium supernatant were collected for TCID_50_ (F, H). \**p* < 0.05, \*\**p* < 0.01, \*\*\**p* < 0.001, \*\*\*\**p* < 0.0001.

### ARV infection induced changes in cellular lipid metabolism

To better understand the mechanism underpinning how ARV infection induced LD formation in infected cells, we performed lipidomics analyses in ARV-infected LMH cells to characterize changes in lipid metabolism compared to mock-infected controls (Figure 5, and Figure S5). Briefly, cells were subjected to untargeted liquid chromatography-mass spectrometry (LC-MS), which revealed that ARV-infected cells had 25-fold higher total lipid content than mock-infected cells. Moreover, the proportion of phosphatidylcholines (PC) in ARV-infected cells was 57.93% compared to only 23.49% in mock-infected cells, while triglycerides (TG) were only 3.41% in ARV-infected cells compared to 51.21% in mock infected cells (Figure 5). Furthermore, lipid ontology (LION) analysis showed an enrichment of triglycerols, lipid storage, and lipid droplet pathways in ARV-infected cells compared to mock-infected cells, consistent with increased LD formation observed during ARV infection (Figure S6).

**Figure 5.**
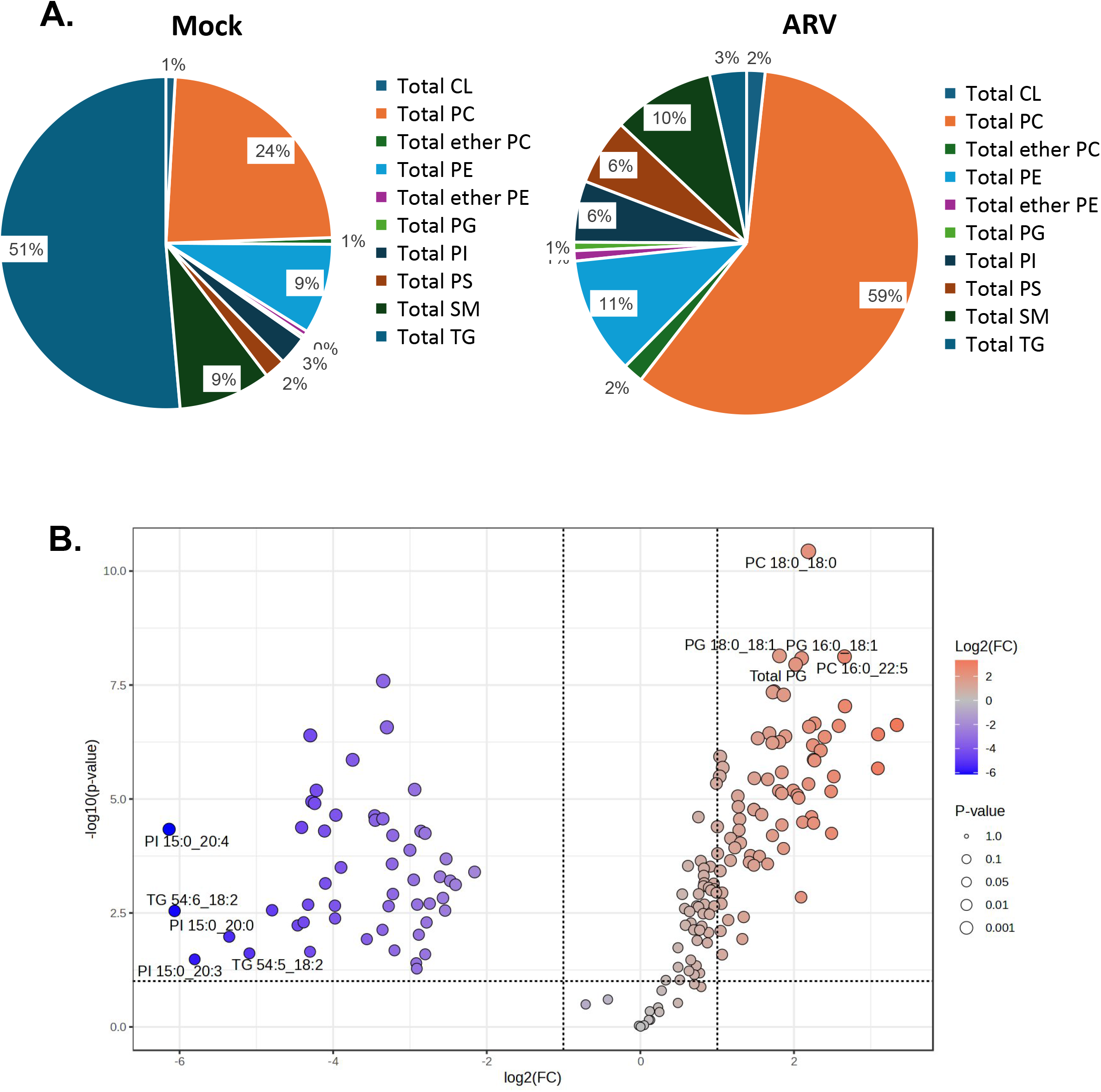
ARV infection led to an increase in total lipid content and phosphotidylcholine fraction. LMH cells were infected with ARV S1133 at an MOI of 3. At 18hpi, cell pellets were harvested and viruses were inactivated through fixation in 95% methanol/water. Cell pellets were sent to the University of Maryland School of Pharmacy lipidmomics core for LC-MS. Lipid counts were normalized to total lipid content for ARV-infected and mock-infected samples, and proportions of lipid classes were calculated (A). Differentially expressed lipids were identified using MetaboAnalyst (v6.0) (B).

Quantification of unsaturated FAs (UFAs) revealed a lower total polyunsaturated FA (PUFA) content in ARV-infected cells, whereas monounsaturated FA (MUFA) content did not significantly change (Figure 6A). Furthermore, there was a decreased total cellular PUFA:MUFA ratio (Figure 6B) that was also observed the PC, phosphatidylserine (PS), and phosphatidylinositol (PI) compartments in ARV-infected cells (Figure 6C). Since PC, PS, and PI are found in cellular membranes^12^, these data suggested a decrease in PUFA:MUFA ratio in cell membranes upon ARV infection. Furthermore, ARV-infected cells had slightly higher PUFA:MUFA ratio in the TG compartment compared to mock infection (Figure 6C).

**Figure 6.**
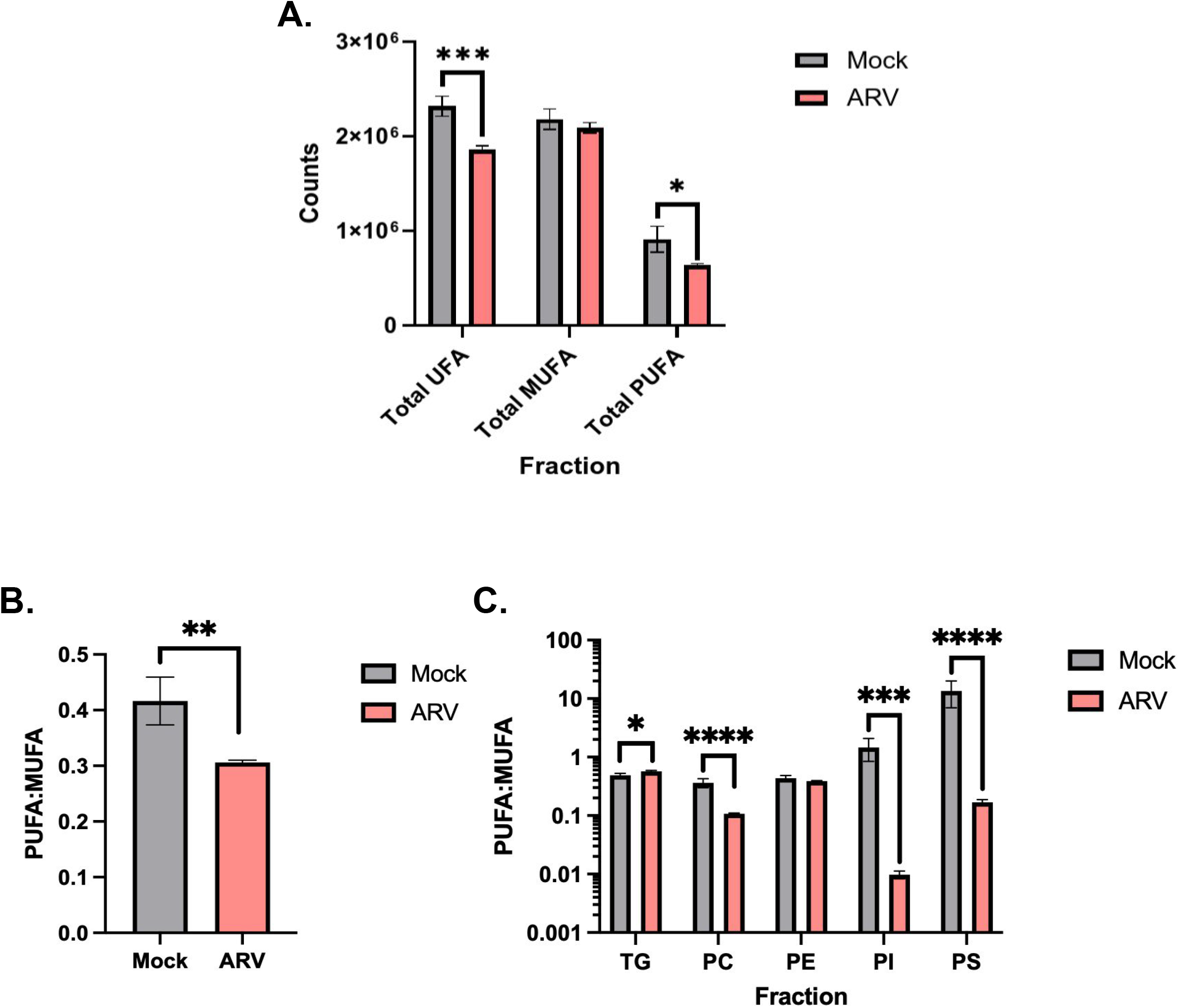
ARV infection led to a decrease in total PUFA content, with depletion of PUFA-PL and relative enrichment of PUFAs in the triglyceride fraction. (A) Counts of total UFAs, MUFAs, and PUFAs were calculated for mock-infected and ARV-infected samples. PUFA:MUFA ratios were calculated in all fractions (B) or separately in different lipid species (C). \**p* < 0.05, \*\**p* < 0.01, \*\*\**p* < 0.001, \*\*\*\**p* < 0.0001.

Next, we performed pathway enrichment analysis from bulk RNA sequencing data. This revealed a decrease in the expression of genes involved in the biosynthesis of UFAs (Figure 7), consistent with our observations of a lower total PUFA content during ARV infection. Moreover, differentially expressed gene (DEG) analysis showed a decreased expression of genes involved in the synthesis of long-chain PUFAs in ARV-infected cells, including elongation of very long-chain FA (*ELOVL*) family and FA desaturase 1 and 2 (*FADS1* and *FADS2*)^13,14^ (Figure 8A). Decreased expression of *ELOVL2* following ARV infection was also confirmed by RT-qPCR (Figure 8B). Taken together, the results of lipidomics and transcriptomics analyses were consistent with ARV infection modulating cellular lipid metabolism, notably causing an increase in total PCs and a shift in MUFA and PUFA metabolism, with decreased PUFA production and depletion of PUFA-phospholipids (PLs) relative to MUFA-PLs, while the proportion of PUFAs in TGs increased.

**Figure 7.**
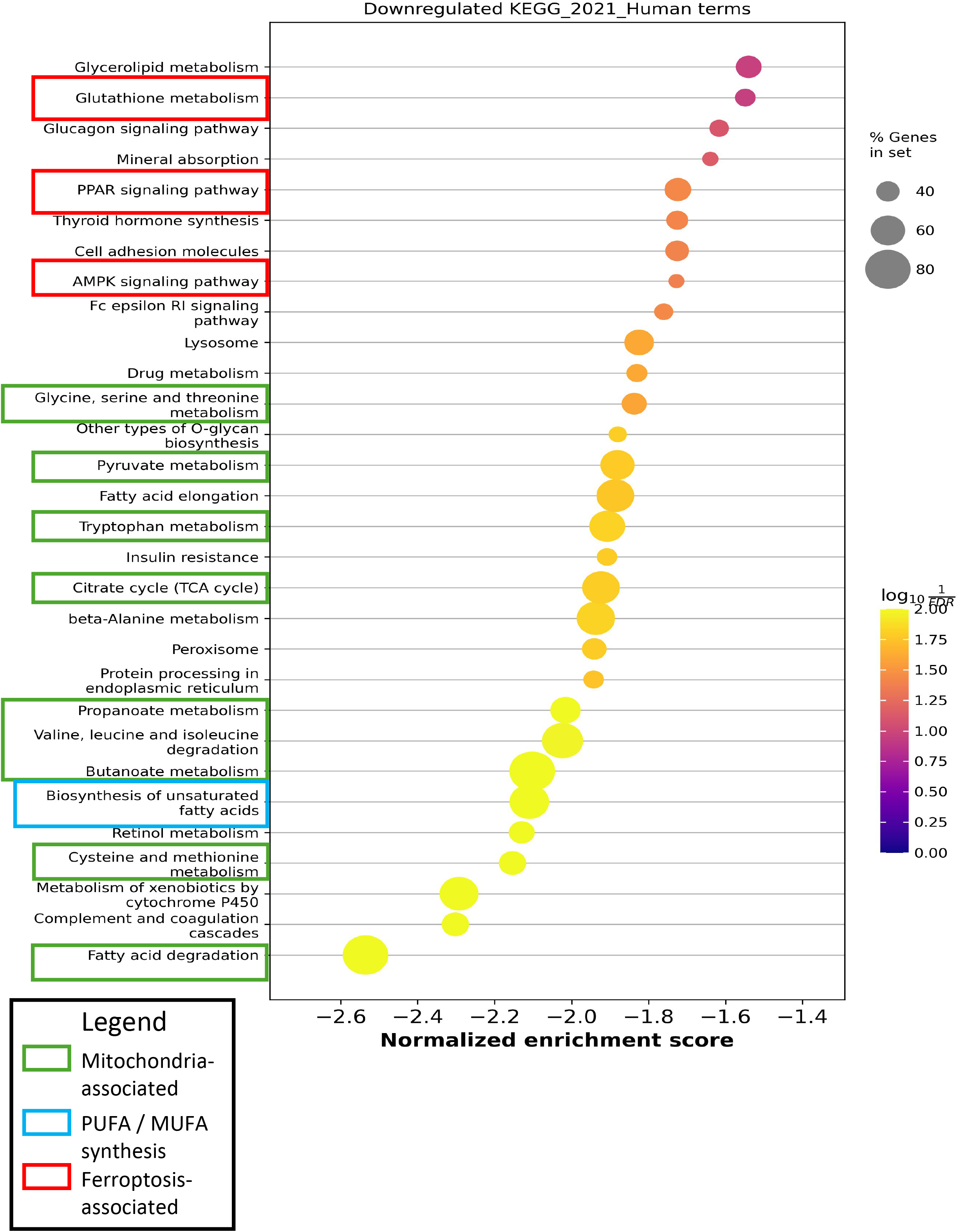
Pathway analysis of bulk-RNASeq dataset revealed that ARV-infected cells had lower expression of PUFA synthesis pathway genes, with changes in mitochondrial metabolism and ferroptosis pathways. LMH cells were infected with ARV S1133 at an MOI of 3. At 18hpi, cells were lysed and RNA was collected and extracted for bulk RNA-seq. Gene set enrichment analysis was performed using GSEApy (v1.3.1). Differentially expressed genes were identified using pyDESeq2 (v0.5.4)

**Figure 8.**
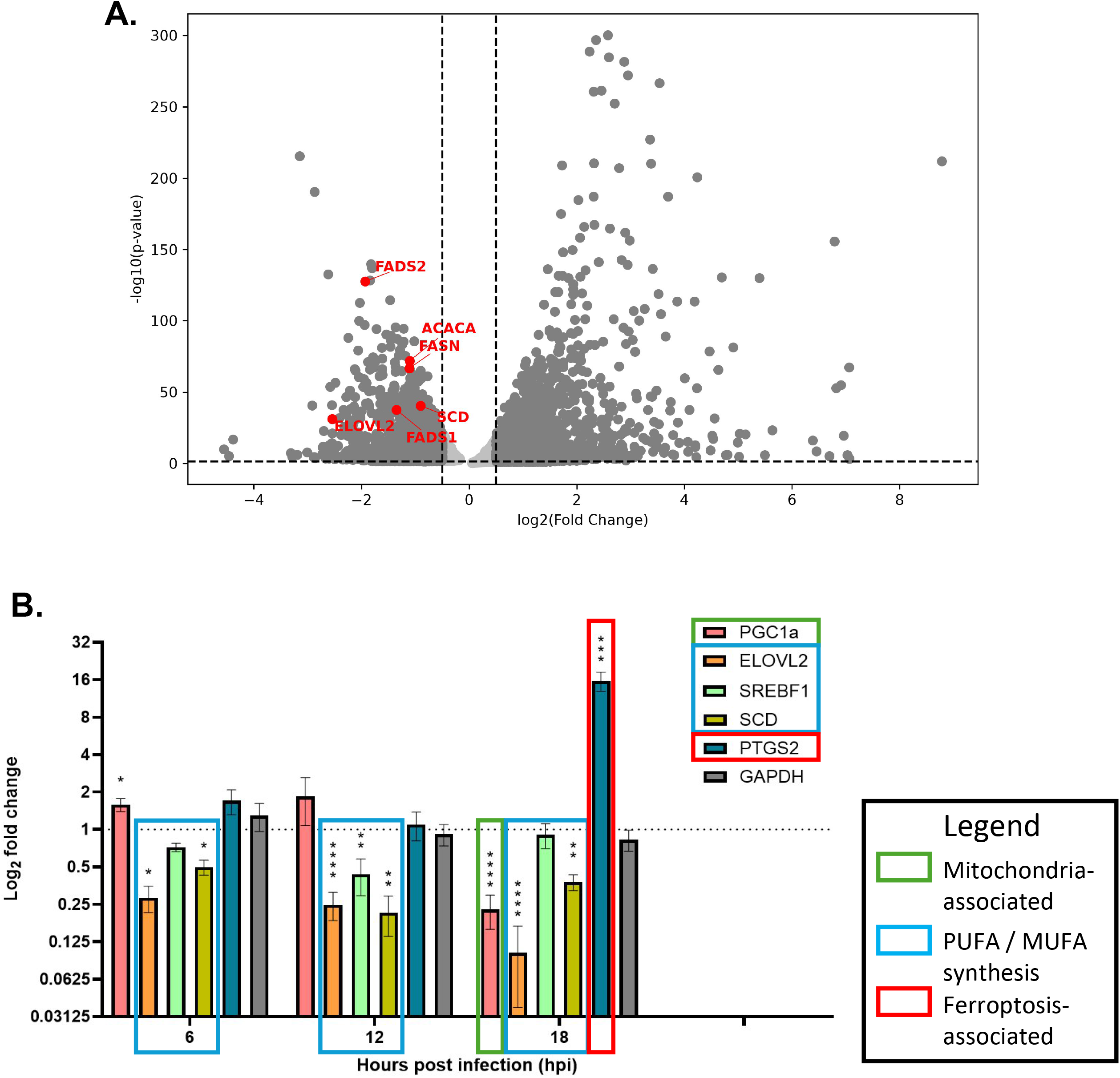
Volcano plot of bulk RNA-Seq dataset, and RT-qPCR confirmation of gene expression. Volcano plot of the bulk RNA-seq dataset (A). Confirmation of bulk RNA-seq results for *PGC1α*, *ELOVL2*, *SREBF1*, *SCD*, and *PTGS2* was performed through RT-qPCR (C). \**p* < 0.05, \*\**p* < 0.01, \*\*\**p* < 0.001, \*\*\*\**p* < 0.0001.

### ARV induced ferroptosis in infected cells, which was proviral at late time points

Cellular membranes with a high PUFA-PL composition are known to be more prone to lipid peroxidation during cellular oxidative stress, which can trigger ferroptotic pathways leading to cell death^2,15,16^. As several viruses, including the dsRNA viruses RV and novel duck orthoreovirus (NDRV), induce ferroptosis following infection^17–19^, we hypothesized that ARV infection also triggered ferroptosis, and that depletion of PUFA-PLs relative to MUFA-PLs may be a means by which cell death is restrained, to facilitate viral replication. To investigate this further, we evaluated whether ARV infection induced cellular oxidative stress, lipid peroxidation, and ferroptosis.

First, we interrogated our bulk RNA-seq pathway enrichment analysis and found that ARV infection led to a downregulation of pathways involved in pyruvate metabolism, the TCA cycle, amino acid metabolism, and FA degradation. This pattern is consistent with mitochondrial dysfunction following ARV infection^20–22^ (Figure 7). RT-qPCR data also showed a significant downregulation of PPARγ coactivator-1α (PGC-1α) expression during ARV infection (Figure 8B). PGC-1α is a transcriptional regulator of genes essential in mitochondrial biogenesis, including mitochondrial antioxidant genes, such as manganese superoxide dismutase^23,24^ . As mitochondrial dysfunction has been shown to increase reactive oxygen species (ROS), these transcriptional data were consistent with our hypothesis that ARV infection could lead to increased cellular oxidative stress^20,22^. Furthermore, glutathione signaling and peroxisome proliferator-activated receptor (PPAR) signaling pathways were also downregulated in ARV-infected cells (Figure 7). Glutathione and glutathione peroxidase are free-radical scavengers critical for antioxidant defense, and decreased glutathione metabolism could be suggestive of lower antioxidant capacity in infected cells^9,15,17,25^. The PPAR pathway regulates antioxidant gene expression, including the genes *PGC-1α*, *GPX4*, and *NRF2*, further indicating decreased antioxidant capacity upon ARV infection^26,27^. Moreover, expression of *PTGS2* was upregulated following ARV infection at 18hpi (Figure 8B). PTGS2 (prostaglandin-endoperoxide synthase 2), also known as COX-2, is a common marker of ferroptosis activation^15,28,29^. Taken together, our transcriptional data indicated possible mitochondrial dysfunction associated with ROS accumulation, decreased antioxidant pathway gene expression, and an increased expression of the ferroptosis marker *PTGS2*, pointing to possible ferroptosis activation during ARV infection.

Next, we measured cellular ROS levels using the Amplex Red assay, with erastin, a ferroptosis inducer, as a positive control. ARV-infection led to a gradual increase in ROS levels, reaching statistical significance at 24hpi (Figure 9A). As ROS-mediated lipid peroxidation is a hallmark of ferroptosis, we next measured lipid peroxidation using BODIPY 581/591 C11, and found that ARV infection led to a significant increase in lipid peroxidation relative to mock-infected cells at 18hpi (Figure 9B, red bars). Finally, as ferroptosis is dependent on intracellular Fe^2+^ , which participates in the Fenton reaction to induce free radical formation and lipid peroxidation^9,13^, we pretreated cells for 6 hours with deferoxamine (DFO), an iron chelator known to inhibit iron-dependent lipid peroxidation and ferroptosis^15,30,31^. DFO pre-treatment partially decreased ARV-induced lipid peroxidation (Figure 9B, green bars). Taken together, these data demonstrate that ARV infection increased oxidative stress and lipid peroxidation that was partially iron-dependent during late infection, consistent with ARV inducing ferroptosis.

**Figure 9.**
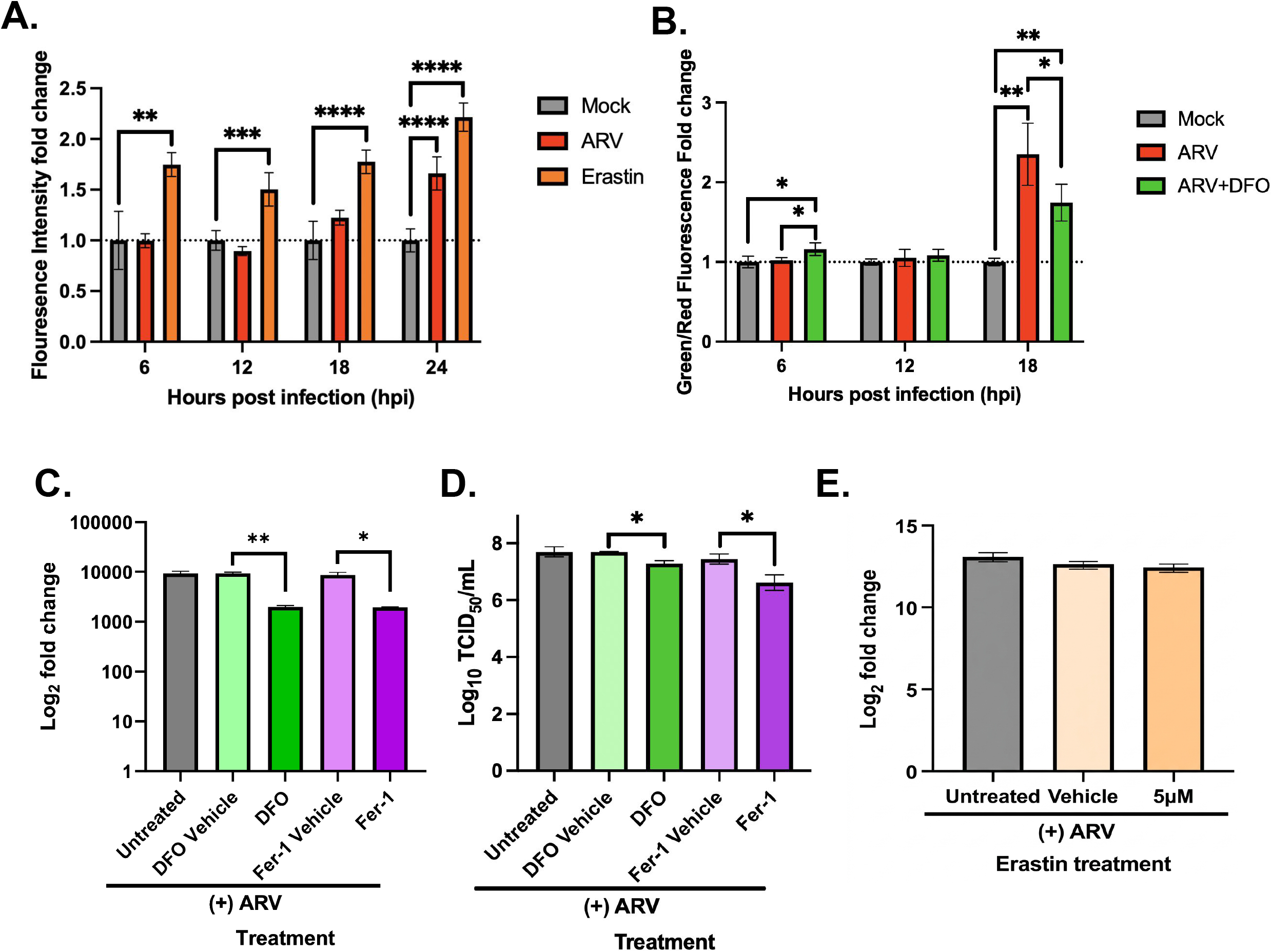
ARV induced ferroptosis in infected cells, and ferroptosis was proviral at later timepoints. (A) LMH cells were infected with ARV S1133 at an MOI of 3, with an erastin-treated uninfected positive control. Cell culture supernatant were collected for Amplex Red assay to quantify ROS released from cells. (B) LMH cells were pre-treated with DFO 6 hours before infection with ARV S1133 at an MOI of 3. BODIPY 581/591 C11 was used to measure lipid peroxidation. (C-D) LMH cells were pre-treated with DFO for 6 hours before infection with ARV S1133 at an MOI of 3. At 18hpi, cells were lysed for quantification of viral transcripts, and culture supernatant were collected for TCID_50_ analysis. (E) LMH cells were pre-treated with erastin 6 hours before infection with ARV S1133 at an MOI of 3. At 18hpi, cell culture supernatant were collected for TCID_50_ analysis. \**p* < 0.05, \*\**p* < 0.01, \*\*\**p* < 0.001, \*\*\*\**p* < 0.0001.

Finally, we sought to determine whether ferroptosis was pro or anti-viral in ARV infection. Inhibition of ferroptosis using DFO or ferrostatin-1 (Fer-1), used at concentrations that did not significantly affect cell viability (Figure S7), led to a significant decrease in ARV mRNA expression (Figure 9C) and viral titer by TCID_50_ (Figure 9D) in LMH cells, suggesting that ferroptosis was proviral for ARV. However, the effect was small, and ferroptosis induction through erastin treatment had no statistically significant effect on ARV replication (Figure 9E). Ferroptosis is thought to favor the spread of viruses to neighboring cells, and in this regard can be considered pro-viral in many viruses^9,13,17,18,32^. Our data suggest ARV-induced ferroptosis during late infection favored viral replication, although the effect was small.

### The ARV p10 protein increased LD formation in transfected cells

To determine the ARV proteins that drive the changes in lipid metabolism and LD formation, we transfected LMH cells with plasmids that expressed each of the ten genome segments individually. Transfection with genome segments S1 and L1 led to the most statistically significant increases in LD number compared to mock-transfected cells, with S1 leading to the greatest increase in average number of LDs per nucleus, while S3, M1, and M2 also increased LD formation to a lesser extent (Figure 10A). S1 encodes three viral proteins: p10, p17, and sigmaC (σC)^10^. We therefore transfected LMH cells with plasmids expressing these proteins individually, fused to an HA tag (HA-p10, HA-p17 and HA-σC), and found that HA-p10 induced a statistically significant increase in the average number of LDs per nucleus compared to an empty vector control (Figure 10B). Taken together, these data suggest that while multiple ARV proteins induce an upregulation of LDs, p10 is a major driver.

**Figure 10.**
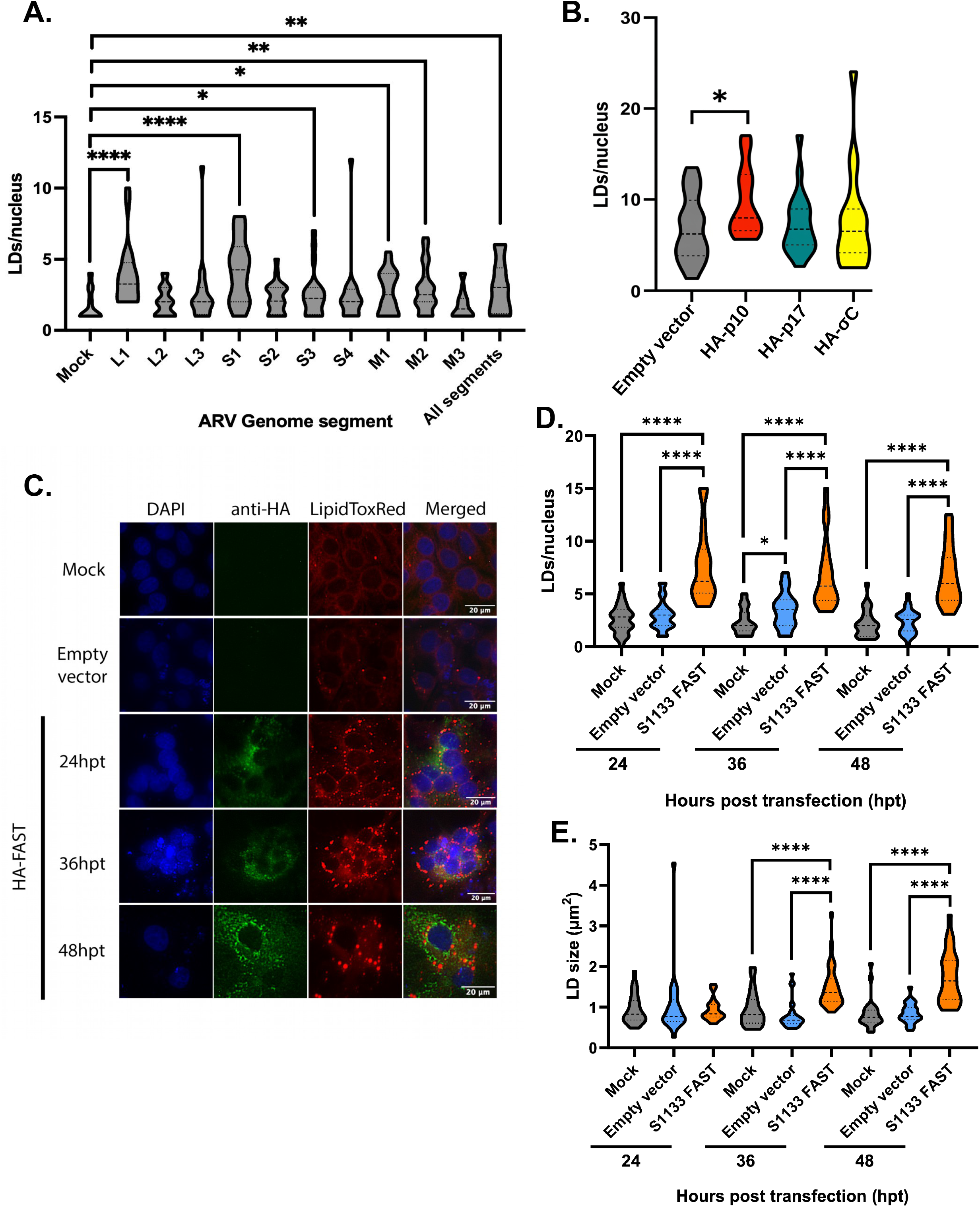
The ARV p10-FAST protein increased LD formation. LMH cells were transfected with individual ARV viral genome segments (A) or plasmids encoding proteins expressed from the S1 gene segment conjugated to HA (B-E). At 24hpt, cells were fixed in 4% PFA and stained with LipidTOX Red for LD quantification. \**p* < 0.05, \*\**p* < 0.01, \*\*\**p* < 0.001, \*\*\*\**p* < 0.0001.

The ARV p10 protein has been identified as a fusion-associated small transmembrane protein (FAST), which induces plasma membrane fusion between neighboring cells, leading to syncytium formation^10,33,34^. Syncytium formation facilitates ARV viral spread between cells^33–35^ and is associated with considerable cellular membrane remodeling^36^. We therefore hypothesized that p10-FAST may be one of the viral proteins inducing changes in lipid metabolism. To investigate this, LMH cells were transfected with HA-p10/FAST, which successfully led to syncytium formation (Figure 10C), and an increase in LD number at 24, 36, and 48 hours post-transfection (hpt) (Figure 10D) as well as increased LD size at 36 and 48hpt (Figure 10E). HA-p10/FAST transfection also led to decreased *ELOVL2* gene expression at 24hpt and progressively increasing *PTGS2* expression from 24-48hpt by RT-qPCR, suggesting decreased PUFA synthesis and increased expression of ferroptosis marker genes (Figure 11A). Consistent with this, HA-p10/FAST transfection led to increased ROS measured by Amplex Red at 24 and 48hpt (Figure 11B), and an increase in lipid peroxidation measured by BODIPY 581/591 C11 at 24 and 36hpt (Figure 11C). Overall, our data suggest that the ARV p10/FAST protein contributed to the induction of ferroptosis, modulation of lipid metabolism, and upregulation of LD formation that we observed following ARV infection.

**Figure 11.**
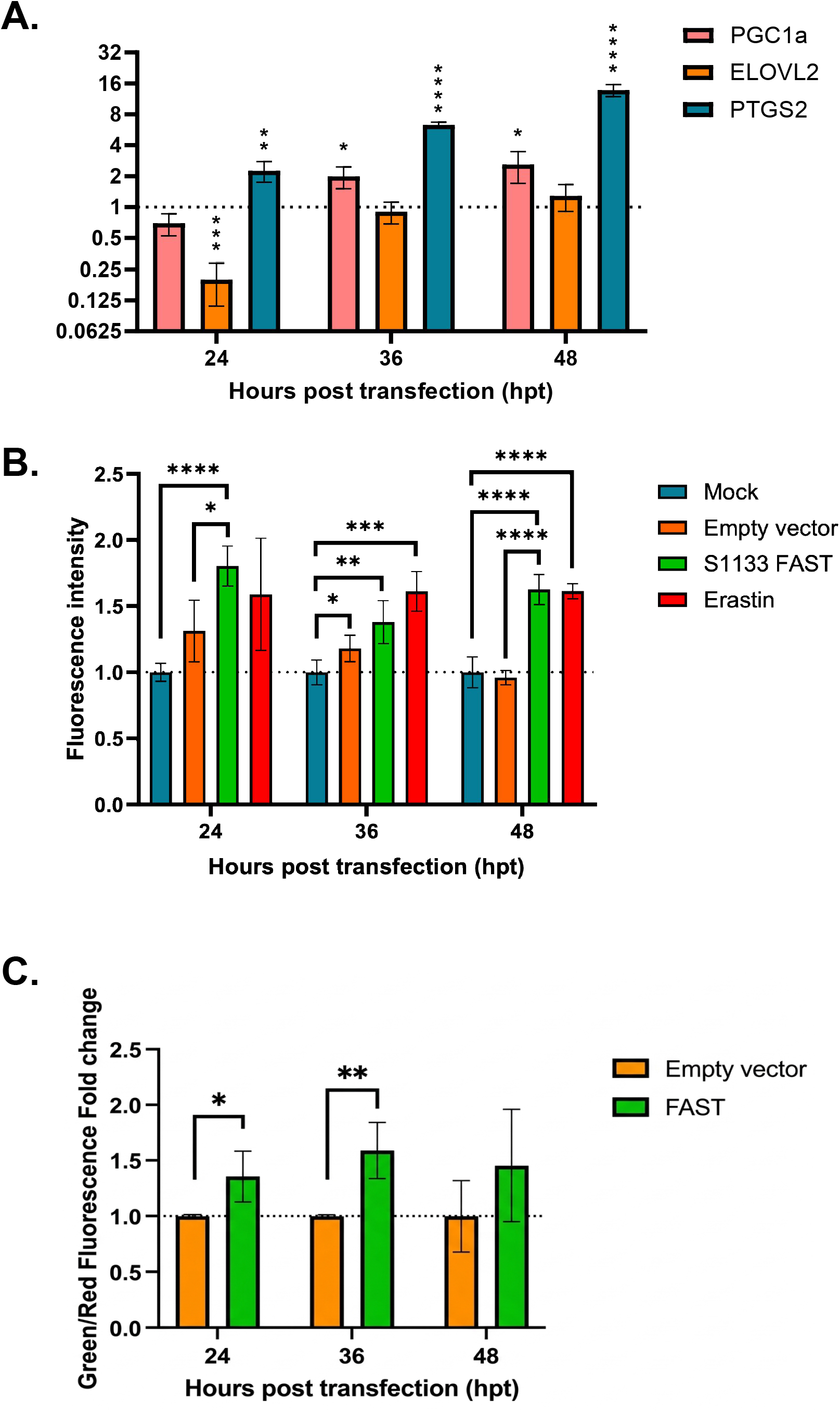
p10-FAST increased *PTGS2* expression and ROS and lipid peroxidation levels in transfected cells. (A) LMH cells were transfected with HA-FAST. Cells were lysed at 24hpt for RT-qPCR quantification of *ELOVL2* and *PTGS2*. (B-C) LMH cells were transfected with HA-FAST or an empty vector control. At 24hpt, supernatants were collected for Amplex Red quantification of ROS (B) or BODIPY 581/591 C11 quantification of lipid peroxidation (C), compared to Mock cells that were untransfected. \**p* < 0.05, \*\**p* < 0.01, \*\*\**p* < 0.001, \*\*\*\**p* < 0.0001.

## Discussion

Members of the *Reoviridae* family are known to utilize LDs during their replication cycle. For example, the replicative compartments of RV and GCRV colocalize with LDs, and these viruses are thought to utilize LDs as a scaffold to nucleate the formation of the compartments^3,5,6^. Furthermore, during mammalian orthoreovirus infection, the outer capsid proteins bind LDs and are redirected away from less mature factories towards more mature factories, to allow sufficient viral replication to proceed prior to virion assembly later in the replication cycle^6^. In contrast, less is known regarding the role of LDs in the replication cycle of the *Birnaviridae* family. In this study, our data demonstrated that the birnavirus IBDV had a less significant effect on LD upregulation compared to the reovirus ARV, demonstrating that the two viruses had a different effect on pathways leading to LD formation. Moreover, neither inhibition nor overexpression of LDs had a significant effect on IBDV replication. These data are consistent with a recent report from Frontini-Lopéz *et al*., who also demonstrated that LDs did not play a significant role in IBDV replication in QM7 cells^37^. We have previously shown that IBDV forms virus factories through liquid-liquid phase separation (LLPS), and that viral protein (VP)3 is necessary for driving this process^38,39^. VP3 is known to interact with the cytoplasmic leaflet of endosomal membranes containing Phosphatidylinositol 3-phosphate (PI3P)^40^, and it is possible that this interaction nucleates the formation of the factories. Based on the data from this current study, it is unlikely that IBDV also utilizes LDs for factory nucleation. This expands our knowledge of IBDV-cell interactions. In addition, even though ARV led to an upregulation of LDs in infected cells, we found that neither inhibition nor overexpression of LDs affected viral replication, and that the ARV VFs did not colocalize with the LDs, suggesting that they do not play a role in the nucleation of the replicative compartments of ARV, in contrast to RV and GCRV. Therefore, our data expand our knowledge of how the *Reoviridae* interact with host cells.

Members of the *Reoviridae* are known to trigger pathways that lead to apoptosis, necroptosis, pyroptosis, autophay, and ferroptosis^17,41–43^. However, there is a paucity of information regrading the cell death pathways triggered by ARV. Previous work has demonstrated the virus can induce apoptosis and autophagy^36,44^, but other cell death pathways, including ferroptosis, are poorly described. Recently, novel duck reovirus (NDRV) was shown to induce ferroptosis^18,19^, so our data therefore expands our knowledge of the cell death pathways triggered by ARV, providing new information regarding the interaction of ARV with host cells. Triggering cell death pathways is thought to favor the spread of viruses to neighboring cells^9,17,18,32,35,41^. However, reoviruses are also known to restrain premature cell death to prolong the longevity of the infected cell, to maximize viral replication prior to triggering cell death later in infection^45,46^. For example, ARV stimulates cell survival pathways early in infection through the activation of PI3K/Akt/NF-κB and STAT3, thus antagonizing premature apoptosis^47^. Ferroptosis can be restrained by downregulating phospholipid PUFAs, as cellular membranes with a high PUFA-PL composition are more prone to lipid peroxidation during cellular oxidative stress, which can trigger ferroptosis^2,15,16^. This can be accomplished by reducing PUFA synthesis in cells, and by rerouting PUFAs away from membrane phospholipids and sequestering them in LDs^2^. Moreover, Rabies virus and encephalomyocarditis virus are known to utilize LD formation to protect cells against arachidonic acid-induced ferroptosis in favor of viral replication^48^. The changes in lipid metabolism we observed following ARV infection included a lower total PUFA content, decreased PUFA:MUFA ratio within the PC, PS, and PI compartments, and an increased PUFA:MUFA ratio in the TG compartment, together with enrichment of lipid storage and lipid droplet pathways. We therefore propose that this modulation in lipid metabolism is an attempt to restrain ARV-induced ferroptosis.

ARV is also known to induce the formation of syncytia, which favors genome mixing and cell-to-cell spread. Syncytium formation involves considerable remodeling of the plasma membrane, and excessive syncytia formation can damage cell membranes, compromise the structural integrity of the cell, and trigger apoptosis^33–36^. We therefore reasoned that the modulation of lipid metabolism may also be a means by which premature cell death induced by excessive syncytium formation might be restrained. Moreover, by decreasing the PUFA:MUFA ratio in phospholipids, there may be an increase in lipid packing in cellular membranes and an increase in rigidity, or resistance to bending or deformation^49–51^, further restraining syncytium formation. Consistent with this hypothesis, when we transfected cells with plasmids expressing each of the ten ARV genome segments individually, S1 had the greatest increase in average LD number/nucleus, and of proteins encoded by S1 (p10/FAST, p17, and σC), p10/FAST had the greatest increase in LDs/nucleus. Moreover, p10/FAST transfection led to a gene expression profile suggestive of decreased PUFA synthesis and ferroptosis induction, and also increased lipid peroxidation as measured by BODIPY 581/591 C11. These data suggest that, in addition to promoting syncytium formation, the ARV p10/FAST protein contributed to ferroptosis induction and lipid metabolism modulation observed following ARV infection, as well as leading to an upregulation of LD formation. The majority of studies concerning the p10/FAST protein have centered around its fusogenic activity. In this study, we expand our knowledge of the protein by uncovering a consequence of membrane fusion, namely an alteration in cellular lipid metabolism.

These findings are important as ARV causes significant morbidity and mortality in chickens, resulting in substantial losses to the poultry industry^10,52,53^. However, the ARV replicative cycle remains poorly understood. Our results thus shed light on ARV viral-host interactions, specifically on ARV-induced changes in host cell lipid metabolism, oxidative stress, and ferroptosis pathways. In addition to being economically significant poultry pathogens, ARV and IBDV are both potential candidates for use as oncolytic viral therapeutic agents^54–57^. Using ARV for oncolytic viral therapy has distinct advantages as it does not infect non-cancerous mammalian cells efficiently, and it is immunostimulatory, inducing anti-tumor immune responses^54,58–60^. To date, ARV has been shown to replicate efficiently in human hepatocellular carcinoma cells and cause oncolysis through syncytium formation, apoptosis activation, and induction of interferon stimulated genes (ISGs)^54,60^. As other oncolytic viruses, such as Newcastle disease virus and oncolytic adenovirus KD01 also mediate cancer cell killing by inducing ferroptosis^61–64^, it is especially important to study how ARV affects cellular lipid metabolism and ferroptosis induction, to also evaluate its potential oncolytic activity.

Our study is not without limitations. First, we did not explore whether LD induction protects against ferroptosis during ARV infection, or whether preventing LD formation promotes ferroptosis. Previous work has shown that LDs sequester PUFAs away from cellular membranes and mediate balanced release of MUFAs and PUFAs, thus protecting cells from ferroptosis in the absence of infection^2^. In addition, others have shown that downregulation of LD synthesis through DGAT inhibition led to ferroptosis activation and decreased viral replication in cells infected with neuroinvasive viruses such as Rabies virus and encephalomyocarditis virus^48^. It would be interesting to explore the dynamics of LD formation, ferroptosis, and viral replication for ARV. In addition, we did not investigate whether inhibition or upregulation of LD formation affects ARV-induced syncytium formation. As LDs play a significant role in buffering changes in cellular lipid metabolism and regulated release and incorporation of PUFAs and MUFAs in membranes^1,2^, this would be interesting to pursue. In addition, to further our understanding of the ARV replicative cycle, future work could also focus on whether ARV outer capsid proteins associate with LDs and facilitate the spatiotemporal regulation of viral genome amplification and capsid assembly similar to mammalian orthoreovirus^6^.

In summary, we observed a modulation of lipid metabolism in ARV-infected cells that was driven, in part, by the p10/FAST protein, leading to the formation of LDs. The LDs were not involved in the formation of ARV replicative complexes, unlike what has been reported for some other members of the *Reoviridae*, such as RV and GCRV. Instead, we demonstrated that ARV formed syncytia and induced ferroptosis, which can promote viral spread to neighboring cells. However, in excess, both processes could also prematurely lead to cell death, and a delicate balance must therefore be struck to maximize viral replication. We theorize that ARV-mediated modulation of lipid metabolism and LD induction may provide a means by which to achieve this balance, by restraining syncytium-induced cell death and ferroptosis.

## Materials and Methods

### Cell and Viruses

The LMH and DF-1 cell lines (chicken hepatocytes and fibroblasts, respectively) were purchased from American Type Culture Collection (ATCC). Chicken 2D primary gut culture cells were prepared as described previously in Orr et al., 2021^65^. Briefly, intestinal tissue were obtained from White Leghorn chickens (University of Maryland Department of Animal and Avian Sciences), at embryonic day 17 (ED17), and placed in 200IU/mL penicillin/streptomycin. Intestinal tissue was digested in 0.2mg/mL collagenase at 37°C, then resuspended in 0.1% bovine serum albumin (BSA) and filtered through a 70µm cell strainer. Samples were resuspended into DMEM/F-12 with B-27 supplement (Gibco) and further digested with accutase to form a single cell suspension as described in Orr et al., 2021^65^. Cells were maintained at 37°C with 5% CO_2_. LMH cells were cultured in DME/F-12 (HyClone SH30023) supplemented with 12% premium grade FBS (VWR 97068) and 2% PenStrep (VWR 30-002-Cl). DF-1 cells were cultured in DMEM (Gibco 11995) supplemented with 10% FBS (ThermoFisher Scientific A5256801). The ARV S1133 strain (Charles River) was grown in LMH cells. The IBDV PBG98 strain was a recombinant molecular clone that was rescued and grown in DF-1 cells as previously described^66^. The chicken field strains of ARV, GF2-E and B8-T, were kind gifts from Dr Milos Markis, Aviserve LLC. These viruses had been propagated in embryonated chicken eggs by inoculation into the yolk sac and subsequent harvest of the embryo and homogenization.

### Antibodies

The chicken polyclonal antibody against ARV was purchased from Abcam (AB33986), and the mouse monoclonal antibody against IBDV VP3 was provided by the UK Immunological Toolbox (The Pirbright Institute, UK).

### Immunostaining

Cells were seeded at a density of 1.7 x 10^5^ cells per well onto 12mm glass coverslips in 24-well plates. Prior to fixation, culture media was removed and the cells were washed with phosphate buffered saline (PBS). Cells were fixed in 4% paraformaldehyde in PBS for 15 minutes at room temperature. The cells were then washed in PBS and permeabilized with a 0.1% saponin solution (Thermo Scientific A18820.14). The permeabilization solution was removed, and the cells were incubated in blocking buffer (0.1% saponin in 4% BSA dissolved in PBS) followed by primary antibodies and secondary antibodies (goat anti-chicken and goat anti-mouse antibodies conjugated to Alexa Fluor 488), diluted in blocking buffer. The secondary antibody solution was removed and cells were washed with PBS prior to incubation in a solution of 4′,6-diamidino-2-phenylindole (DAPI) (Thermo Scientific 62248). The DAPI solution was removed, cells were washed, and incubated in HCS LipidTOX Red (ThermoFisher Scientific H34476) diluted 1:1,000 in PBS for 30 minutes at room temperature. The LipidTox Red solution was then removed and cells were washed once with water. The coverslips were mounted onto glass slides with Prolong Glass antifade solution (Invitrogen P36982) and protected from light.

### Imaging

For the infection time course experiments in LMH and DF-1 cells, slides were imaged using the Stellaris confocal microscope (Leica) through the 63x oil objective. For all other experiments, the cells were imaged using a AXIO Observer fluorescence microscope (Zeiss) through a 40x air objective.

### Image Analysis

Detection and quantification of LDs stained with LipidTOX Red was performed using scikit-image (v0.24) in a pipeline developed by Andrew Brodrick (https://github.com/LadInTheLab/objectFinder2). Contours of LDs were enhanced by applying a Sobel filter, then watershed segmentation was performed to segment and label individual LDs for quantification of LD number per nucleus and LD area. Twenty LD quantification data points were acquired from two replicate wells for all experiments.

### Bulk RNA-seq

Cells were infected with ARV at an MOI of 3. At 18hpi, cells were lysed in RLT buffer and RNA was extracted using an RNeasy kit (QIAGEN). RNA was shipped to Novogene, for RNA Sequencing using an Illumina Novoseq platform. Differential gene expression analysis was performed using PyDESeq2 (v0.5.4). Gene set enrichment analysis was performed using GSEApy (v1.3.1).

### Downregulation of LD Formation

TOFA [5-(tetradecyloxy)-2-furoic acid] (GlpBio GC32715), an allosteric inhibitor of acetyl-CoA carboxylase 1 (ACC1), was used to inhibit the rate-limiting step of *de novo* lipogenesis and LD synthesis as previously described^8^. The cytotoxic effect of TOFA was measured using the CyQuant MTT Cell Proliferation Assay kit (ThermoFisher Scientific) to determine the suitable working concentration of TOFA (described below). To eliminate exogenous fatty acids, cells were cultured for one passage in culture media without FBS supplemented with 1mg/mL fatty acid-free BSA (Sigma-Aldrich A8806), then seeded onto 24-well plates in FA-free culture media. TOFA was dissolved in DMSO to a stock concentration of 10mM. The 10mM stock solution was diluted in pre-warmed FA-free cell culture media to the working concentration. Cells were treated with TOFA eight hours before infection by removing media from the well and adding TOFA-treated media, then incubating at 37°C. Cells were infected with ARV or IBDV by adding diluted virus in cell culture media and incubating for one hour at 37°C. After infection, cells were washed once with PBS, and fresh FA-free media containing TOFA was added and cells were incubated at 37°C.

### Upregulation of LD Synthesis

PA and OA were used to upregulate LD synthesis through exogenous fatty acid supplementation. To prepare PA-BSA and OA-BSA conjugates, PA or OA were dissolved in sodium hydroxide pre-warmed to 100°C. The dissolved PA and OA were added to FA-free BSA pre-warmed to 45°C to reach a final concentration of 5mM PA-BSA and OA-BSA stock solutions. The stock solutions were diluted in cell culture media to reach working concentrations of 250µM PA-BSA and 250µM OA-BSA. The working solutions were then mixed at a 2:1 ratio by volume to create a 2:1 OA-BSA:PA-BSA solution. The cells were incubated in the PA, OA, and 2:1 OA:PA solutions at 37°C four hours before infection. Cells were then infected with diluted virus in culture media for one hour at 37°C. After infection, cells were washed once with PBS, and fresh culture media containing PA, OA, or 2:1 OA:PA was added and cells were incubated at 37°C.

### MTT Cell Viability Assay

Cells were plated and a 12mM MTT stock solution was prepared by adding 1mL PBS to a 5mg vial of MTT and stored at 4°C protected from light. The tell culture media was removed from each well, and 100µL of fresh cell culture media without phenol red was added. 10µL of the 12mM MTT stock solution was then added to each well. For the negative control, 100µL cell culture medium and 10µL of MTT stock solution was added to a well without cells. The cells were then incubated at 37°C for two hours. After incubation, 85µL of medium was removed to leave 25µL of medium in each well. 50µL DMSO was then added to each well and mixed by pipetting. Cells were incubated at 37°C for 10 minutes, then mixed again by pipetting. The absorbance at 540nM was measured using the BioTek Synergy HTX Multi-Mode Microplate Reader. Percent cell viability was calculated as follows: 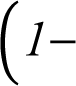 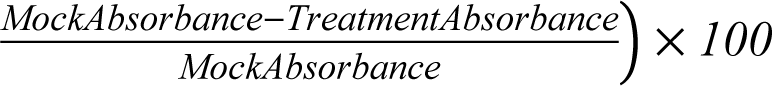. Cell viability of each treatment group was compared to a viability of 100% in the mock-treated group.

### RT-qPCR

Cells were lysed using Buffer RLT (QIAGEN) and kept at -80°C. RNA extraction was performed using the QIAshredder (Cat. No. 79656) and RNeasy Mini (Cat. No. 74106) kits (QIAGEN) as per the manufacturer instructions. For quantifying ARV mRNA levels, the OneStep RT-PCR kit (QIAGEN Cat. No. 210212) was used as per manufacturer instructions. The two-step RT-qPCR assay was used for quantifying IBDV and lipogenesis gene mRNA levels. Briefly, cDNA was synthesized using 5µL extracted RNA template, random primers (Integrated DNA Technologies IDT), and SuperScript III Reverse Transcriptase (Invitrogen 18080093). RT-qPCR was performed using 1:10 diluted cDNA. The Taqman real-time qPCR assay (Applied Biosystems 4366072) was used for quantifying IBDV mRNA levels per the manufacturer instructions. For quantifying lipogenesis gene mRNA levels, the Luna real-time qPCR assay (New England Biolabs M3003) was used per manufacturer instructions. All primers used for RT-qPCR are listed in Table 1. All qPCR reactions were performed using the CFX Connect Real-Time PCR system (Bio-Rad).

**Table 1.**
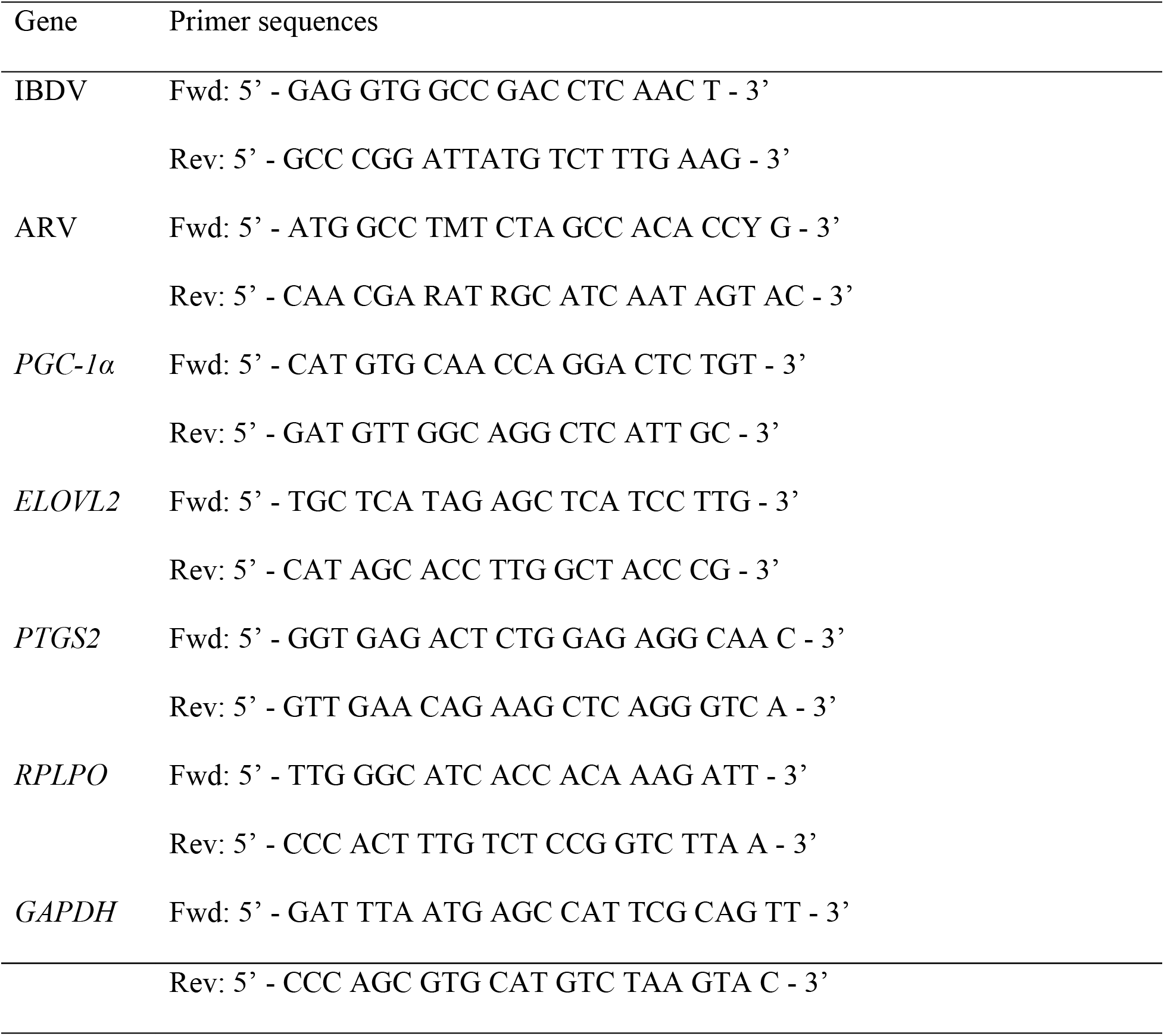
Primer sequences used for RT-qPCR.

### TCID_50_

LMH cells were seeded at a density of 2×10^6^ cells per plate in 96 well-plates with 180µL cell culture medium per well and incubated at 37°C for 24 hours. 20µL of supernatant containing virus was added to quadruplicate wells in the top row and mixed by pipetting. Ten-fold serial dilutions of the virus sample were performed, and cells were incubated for 48-72 hours at 37°C, 5% CO_2_. Infected wells were identified by the appearance of cytopathic effect, and TCID_50_/mL was then calculated.

### Lipidomics analysis

LMH cells were infected with ARV S1133 at an MOI of 3 for 18 hours, whereupon cells were trypsinized and pelleted, and virus was inactivated by resuspending and fixing the pellet in 95% methanol (HPLC grade) in water. Pellets were washed in water, and frozen at -80°C and sent on dry ice to the University of Maryland School of Pharmacy lipidomics core for LC-MS analysis. Differential expression analysis of lipid species was performed with MetaboAnalyst (v6.0)^67^. Lipid ontology analysis was performed using LION^68^.

### Plasmids and Transfection

The coding sequences of the ten segments of the ARV S1133 strain were synthesized by Genscript and cloned into the pSF-CAG-Kan expression vector (Sigma-Aldrich OGS505). The TransIT-LT1 transfection reagent (Mirus Bio MIR2304) was used for all transfection experiments according to manufacturer protocol. A ratio of 3µL TransIT-LT1 per 1µg of DNA was used. Transfection solutions were prepared in Opti-MEM I Reduced-Serum Medium (Gibco 31985070).

### Erastin, deferoxamine, and ferrostatin-1 treatment

Erastin (Sigma-Aldrich 329600) was diluted to a working concentration of 50µM in cell culture medium and added to cells for ferroptosis induction. For ferroptosis inhibition through iron chelation, cells were pre-treated with 200µM of deferoxamine mesylate (Millipore 252750) diluted in cell culture medium six hours before infection or transfection. For ferroptosis inhibition through reduction of ROS species, cells were pre-treated with 5µM of ferrostatin-1 (Sigma-Aldrich SML0583) diluted in cell culture medium six hours before infection or transfection.

### Amplex Red

The Amplex UltraRed assay kit (Invitrogen A36006) was used to measure reactive oxygen species in infected or transfected cells according to manufacturer protocol. HEPES pH 7.0 (ThermoScientific 7365-45-9) was used as the reaction buffer. Cells were cultured and infected or transfected as previously stated. At the analysis timepoint, 50µL of culture media supernatant was transfered to a 96 well black plate (ThermoScientific 137101), and 50µL of 10mM Amplex UltraRed reagent was combined with the culture supernatant. The well plate was covered, protected from light, and shaken on the orbital for 30 minutes at room temperature. A microplate reader was used to measure fluorescence at 530/590nm excitation/emission wavelengths.

### BODIPY 581/591 C11

The BODIPY 581/591 C11 assay kit (Invitrogen D3861) was used for ratiometric measurement of lipid peroxidation according to manufacturer protocol. Cells were seeded in 96 well clear-bottom white well plates (ThermoScientific 165306) and infected or transfected as previously stated. At the analysis timepoint, cell culture media was removed from each well and discarded. 100µL of 10mM BODIPY 581/591 C11 solution was added to each well. Cells were incubated at 37°C for 30 minutes, then washed three times with PBS. Fluorescence was measured with a microplate reader. Fluorescence of oxidized lipids was measured at 488/510nm excitation/emission, while that of reduced lipid lipids was measured at 581/591nm. The ratio of oxidized/reduced lipids was calculated for each well.

### Statistical Analysis

The Mann-Whitney U Test was used to compare LD number and size as well as viral mRNA levels and TCID_50_/mL. The Kruskal-Wallis test with Dunn’s post-hoc pairwise multiple comparison test was used to compare Amplex red, BODIPY 581/591 C11, and PUFA:MUFA ratio measurements between mock-infected and ARV-infected cells. There were a total of 20 data points for LD number and size from two replicate wells for each experiment. Viral RT-qPCR and TCID_50_/mL measurements were taken from three replicates for each experiment. The Kruskal-Wallis Test was used to compare lipogenesis gene mRNA levels. Six replicates from each experiment were used to quantify lipogenesis gene mRNA levels. Statistical analysis and graphs were made with Graphpad Prism 10.

## Acknowledgements

This work was supported by University of Maryland startup funds to A. J. Broadbent. In addition, we acknowledge the Mass Spectrometry Center at the University of Maryland School of Pharmacy for untargeted liquid chromatography-mass spectrometry. The funders had no role in study design, data collection and interpretation, or the decision to submit the work for publication.

---

**Supplementary Figure S1.**
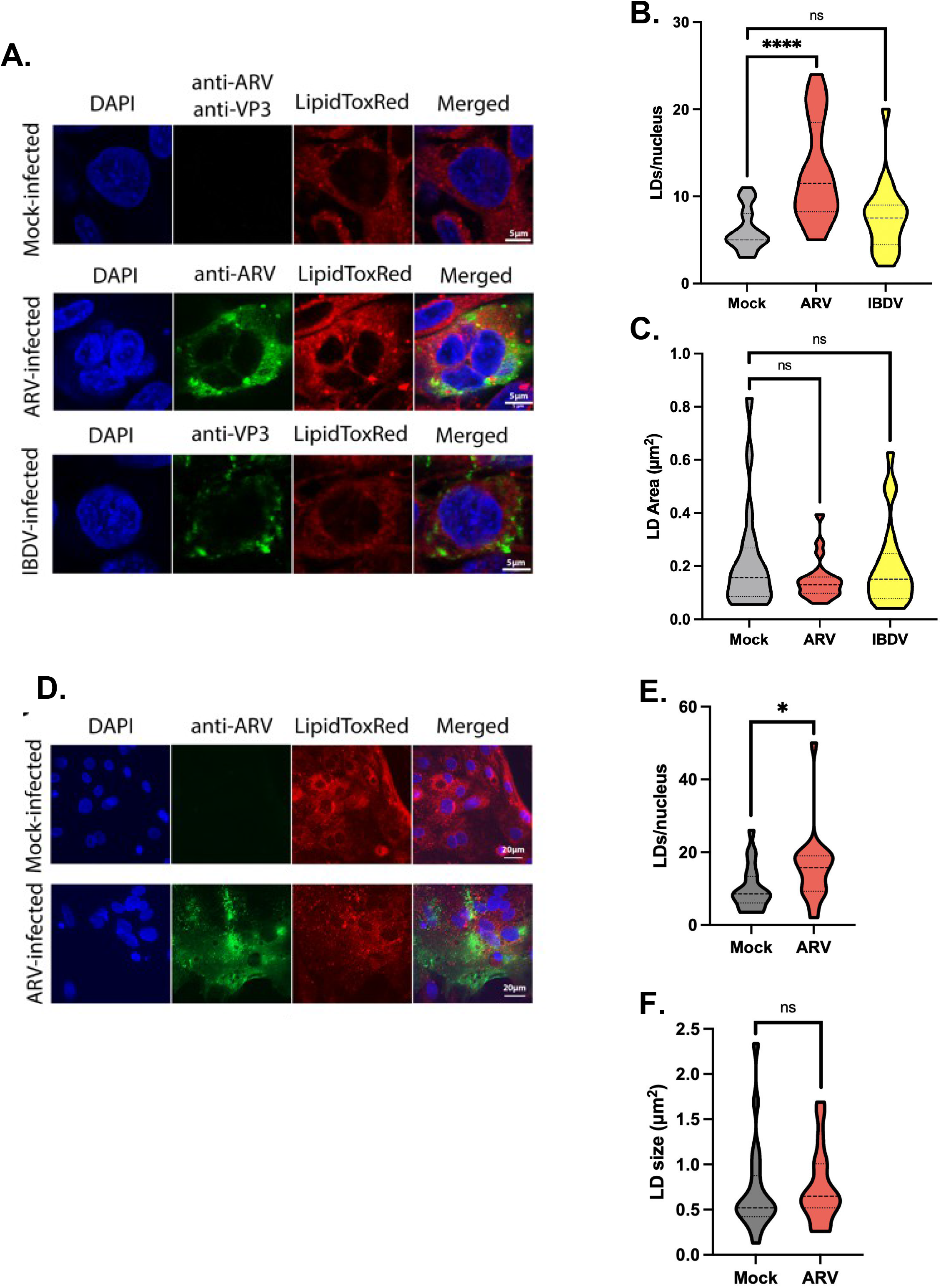
ARV infection increased LD formation in DF-1 and primary gut culture cells. DF-1 cells (A-C) were infected with ARV S1133 or IBDV PBG-98, and chicken primary gut culture cells (D-F) were infected with ARV S1133. Cells were infected at an MOI of 3, fixed at 18hpi and stained with an ARV polyclonal antibody or an IBDV monoclonal antibody raised against VP3, and LipidTOX Red (A, and D), and LDs were quantified (B, C, E, and F). \**p* < 0.05, \*\**p* < 0.01, \*\*\**p* < 0.001, \*\*\*\**p* < 0.0001.

**Supplementary Figure S2.**
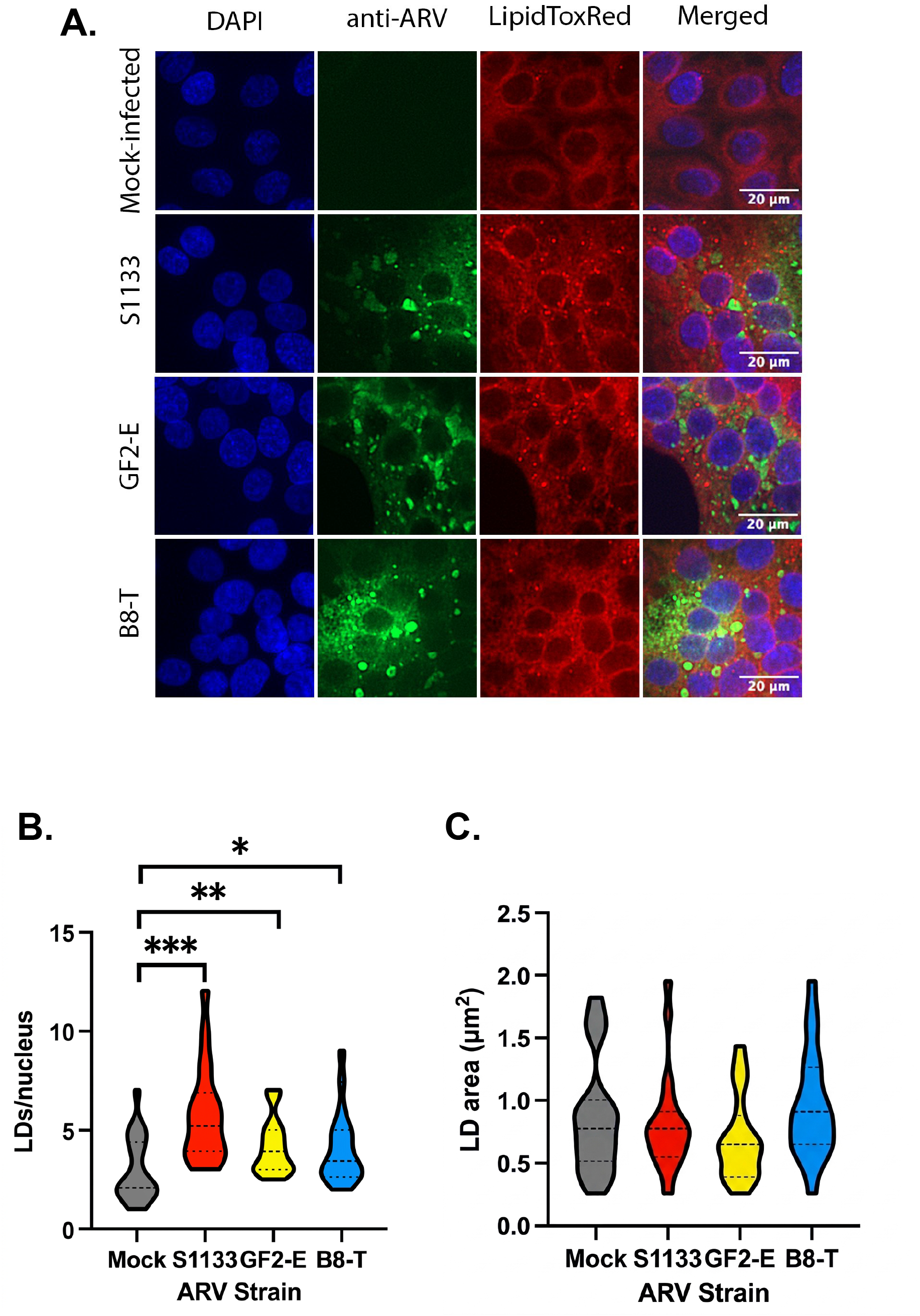
ARV field strains GF2-E and B8-T increased LD formation. LMH cells were infected with the ARV field strains GF2-E and B8-T at an MOI of 1. (A) Cells were fixed at 18hpi and stained with ARV polyclonal antibody. (B-C) LD number and size were quantified. \**p* < 0.05, \*\**p* < 0.01, \*\*\**p* < 0.001, \*\*\*\**p* < 0.0001.

**Supplementary Figure S3.**
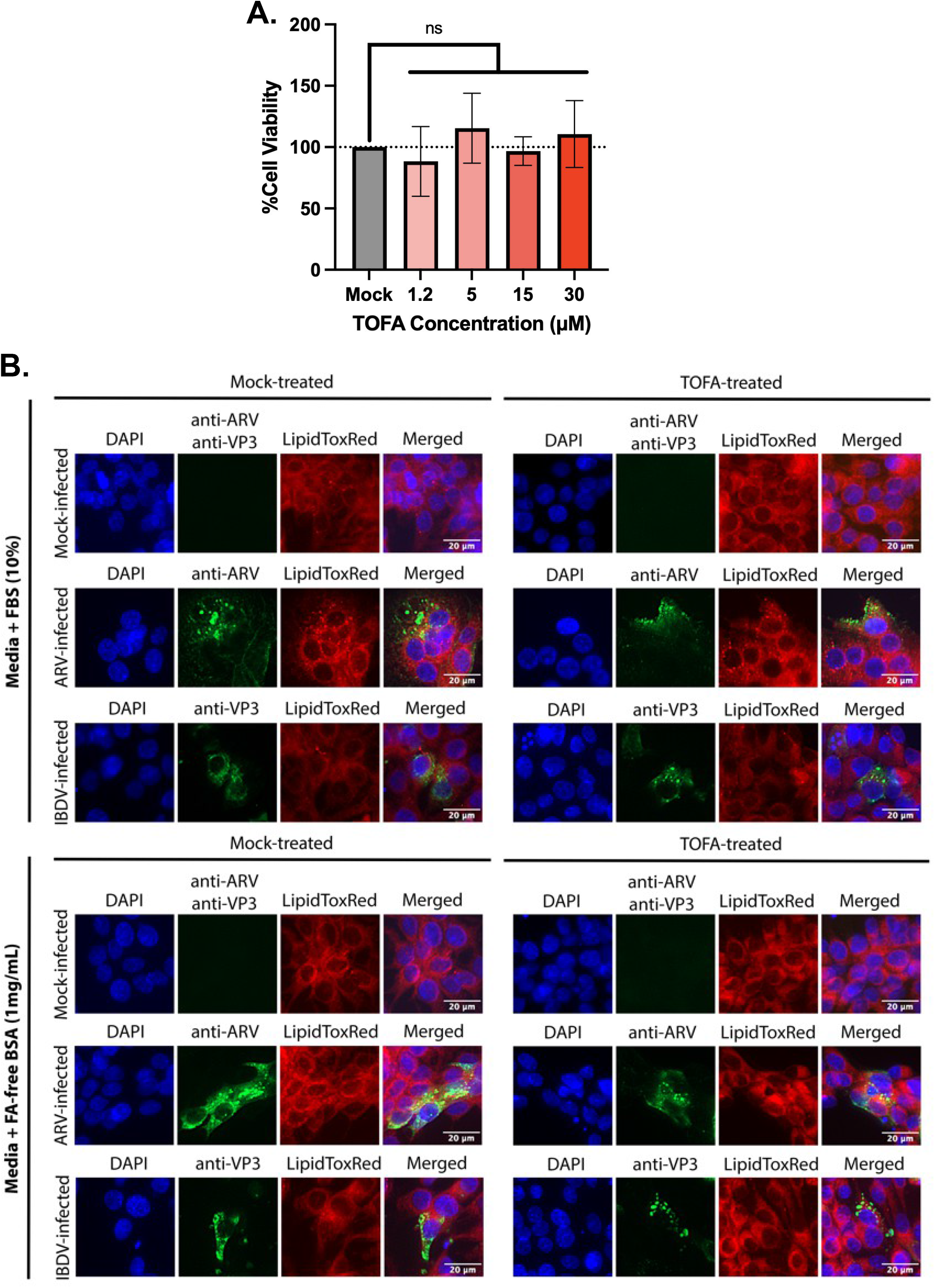
TOFA treatment did not decrease cell viability and successfully inhibited LD formation in FA-free culture conditions. (A) LMH cells were treated with varying concentrations of TOFA or DMSO as mock treatment. The MTT assay was performed 24 hours after treatment to measure cell viability. (B) LMH cells were cultured in either 10% FBS-supplemented or FA-free BSA-supplemented DMEM F-12 for one passage. Cells were then pre-treated with TOFA 6 hours before infection, then infected with ARV S1133 at an MOI of 3. Cells were fixed at 18hpi and stained with ARV polyclonal antibody and LipidTOX Red for LD quantification.

**Supplementary Figure S4.**
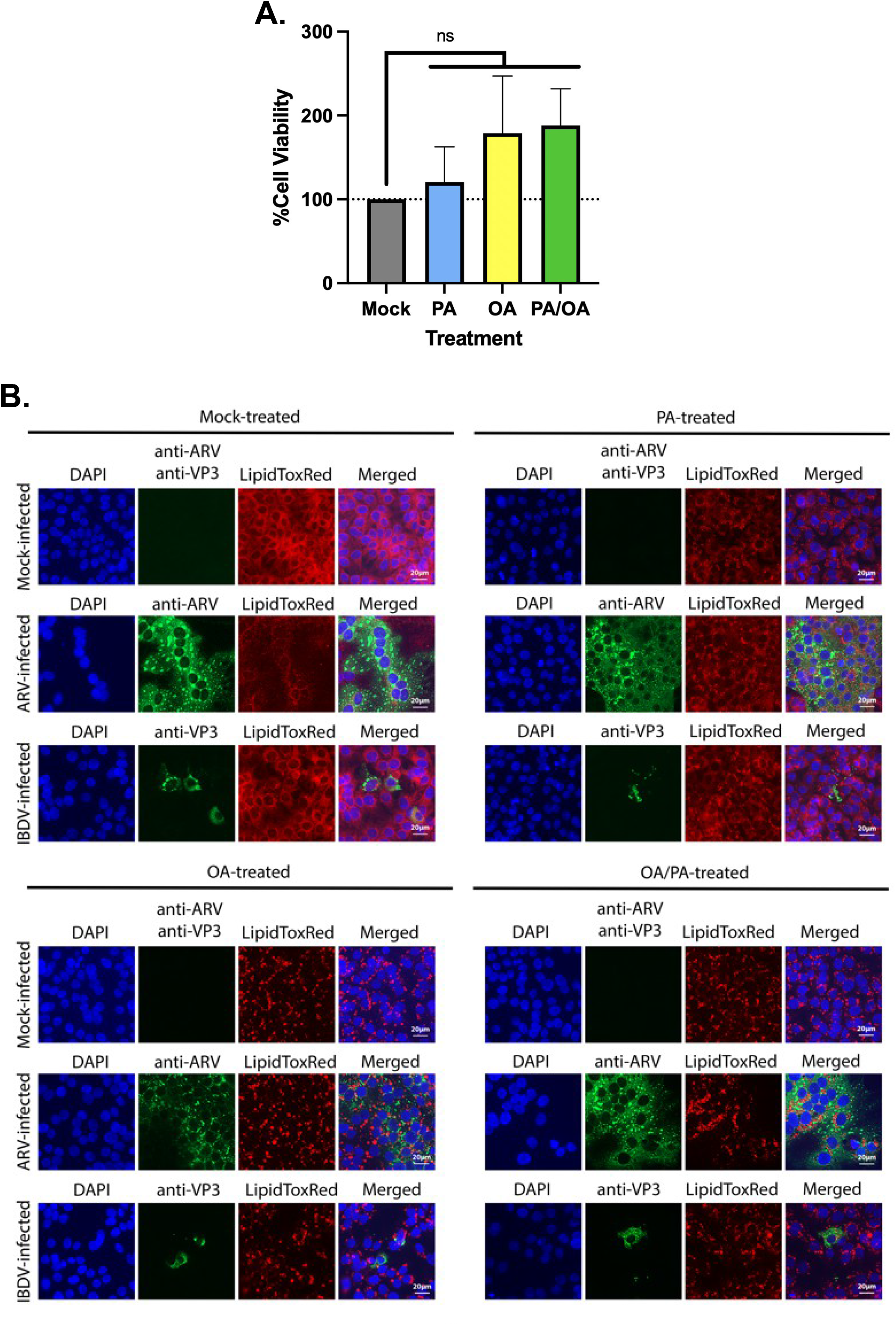
PA and OA treatment did not decrease cell viability and upregulated LD formation. (A) LMH cells were treated with 250µM of PA-BSA, 250µM of OA-BSA, or a 2:1 mixture of OA-BSA:PA-BSA solution. The MTT assay was performed 24 hours after treatment to measure cell viability. (B) LMH cells were pre-treated with the fatty acid solutions 6 hours before infection, then infected with ARV S1133 at an MOI of 3. Cells were fixed at 18hpi and stained with ARV polyclonal antibody and LipidTOX Red for LD quantification.

**Supplementary Figure S5.**
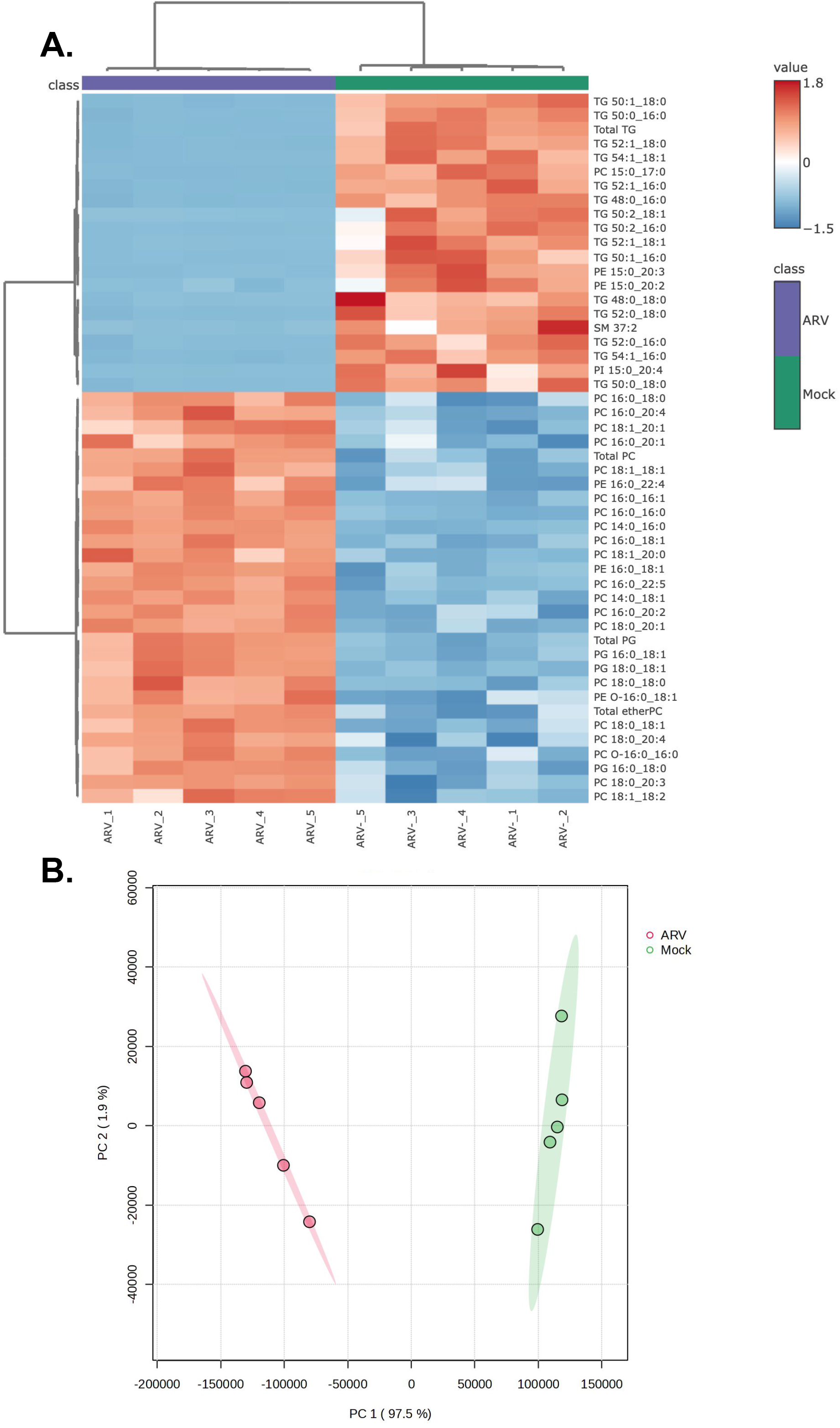
Bulk RNA-seq showed substantial changes in gene expression upon ARV infection. (A-B) LMH cells were infected with ARV S1133 at an MOI of 3 and lysed for RNA extraction at 18hpi for bulk RNA-seq. Differential gene expression heat map (A) and PCA plot (B) show substantial differences in gene expression between mock-infected and ARV-infected cells.

**Supplementary Figure S6.**
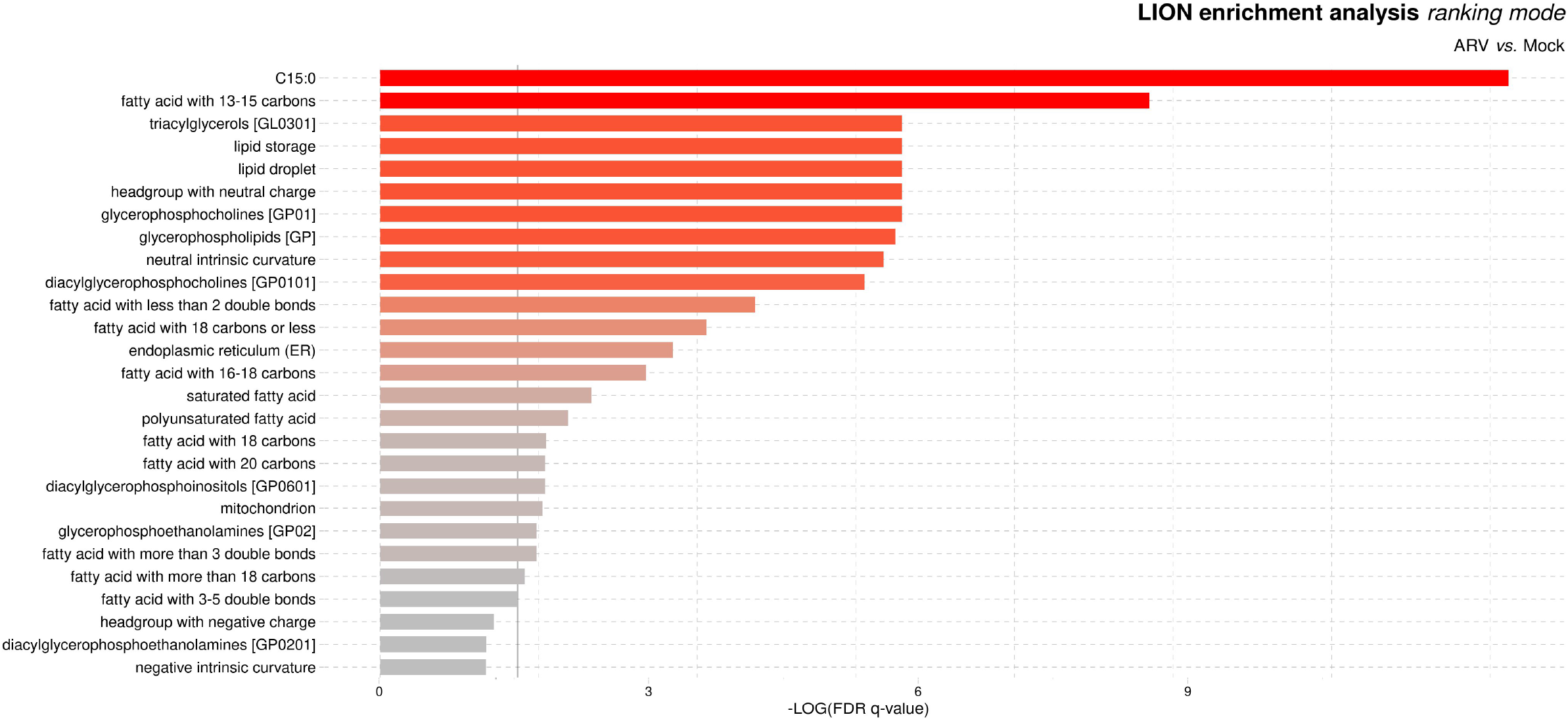
Lipid ontology analysis confirmed induction of lipid droplets upon ARV infection. LMH cells were infected with ARV S1133 at an MOI of 3. Cells were fixed in 95% methanol/5% water to inactivate virus particles, then sent for LC-MS lipidomics analysis. Lipid ontology (LION) analysis showed upregulation of lipid droplet related lipid species.

**Supplementary Figure S7.**
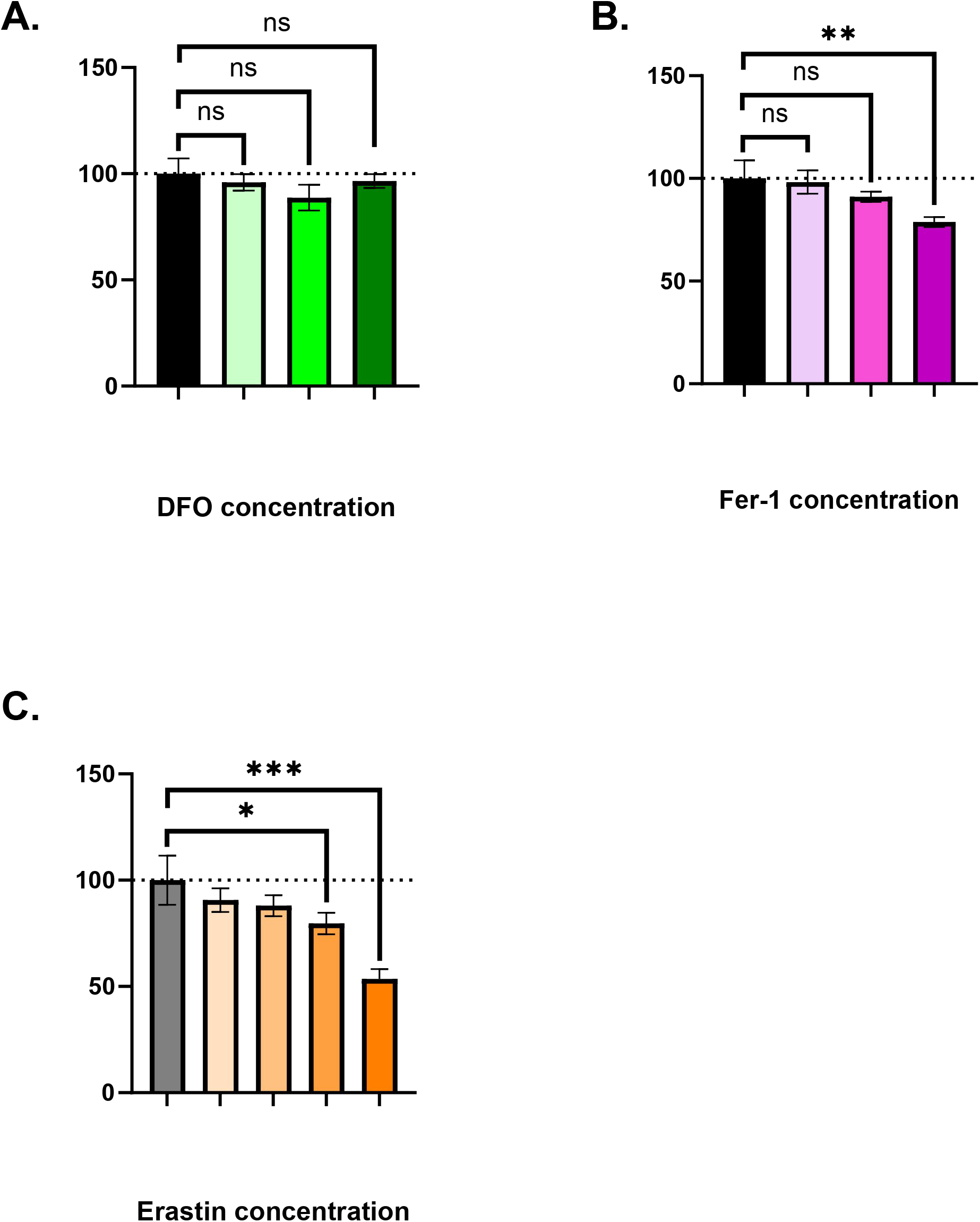
DFO, Fer-1, and erastin treatment’s impacts on cell viability. LMH cells were treated with varying concentrations of DFO (A), Fer-1 (B), or erastin (C) for 24 hours. Cell viability was measured using the MTT assay. \**p* < 0.05, \*\**p* < 0.01, \*\*\**p* < 0.001, \*\*\*\**p* < 0.0001.

